# ForestQC: quality control on genetic variants from next-generation sequencing data using random forest

**DOI:** 10.1101/444828

**Authors:** Jiajin Li, Brandon Jew, Lingyu Zhan, Sungoo Hwang, Giovanni Coppola, Nelson B. Freimer, Jae Hoon Sul

## Abstract

Next-generation sequencing technology (NGS) enables discovery of nearly all genetic variants present in a genome. A subset of these variants, however, may have poor sequencing quality due to limitations in sequencing technology or in variant calling algorithms. In genetic studies that analyze a large number of sequenced individuals, it is critical to detect and remove those variants with poor quality as they may cause spurious findings. In this paper, we present a statistical approach for performing quality control on variants identified from NGS data by combining a traditional filtering approach and a machine learning approach. Our method uses information on sequencing quality such as sequencing depth, genotyping quality, and GC contents to predict whether a certain variant is likely to contain errors. To evaluate our method, we applied it to two whole-genome sequencing datasets where one dataset consists of related individuals from families while the other consists of unrelated individuals. Results indicate that our method outperforms widely used methods for performing quality control on variants such as VQSR of GATK by considerably improving the quality of variants to be included in the analysis. Our approach is also very efficient, and hence can be applied to large sequencing datasets. We conclude that combining a machine learning algorithm trained with sequencing quality information and the filtering approach is an effective approach to perform quality control on genetic variants from sequencing data.

**Author Summary:** Genetic disorders can be caused by many types of genetic mutations, including common and rare single nucleotide variants, structural variants, insertions and deletions. Nowadays, next generation sequencing (NGS) technology allows us to identify various genetic variants that are associated with diseases. However, variants detected by NGS might have poor sequencing quality due to biases and errors in sequencing technologies and analysis tools. Therefore, it is critical to remove variants with low quality, which could cause spurious findings in follow-up analyses. Previously, people applied either hard filters or machine learning models for variant quality control (QC), which failed to filter out those variants accurately. Here, we developed a statistical tool, ForestQC, for variant QC by combining a filtering approach and a machine learning approach. We applied ForestQC to one family-based whole genome sequencing (WGS) dataset and one general case-control WGS dataset, to evaluate our method. Results show that ForestQC outperforms widely used methods for variant QC by considerably improving the quality of variants. Also, ForestQC is very efficient and scalable to large-scale sequencing datasets. Our study indicates that combining filtering approaches and machine learning approaches enables effective variant QC.

## Introduction

Over the past few years, genome-wide association studies (GWAS) have been playing an important role in identifying genetic variations associated with common diseases or complex traits(1, 2). GWAS have found many associations between common variants and human diseases, such as schizophrenia(3), type 2 diabetes(4, 5) and Parkinson’s Disease(6). However, these common variants typically explain only a small fraction of heritability for the complex traits(7, 8). Rare variants are another type of genetic variants that have been considered as an important risk factor for complex traits and common diseases(9–12). With the next generation sequencing (NGS) technology, geneticists may now gain insights into the roles of novel or rare variants. For instance, deep targeted sequencing was applied to discover rare variants associated with inflammatory bowel disease(13). Whole genome sequencing (WGS) has been used to identify rare variants associated with prostate cancer(14), and with whole exome sequencing, studies have also detected rare variants associated with LDL cholesterol(15) and autism(16).

NGS data are not, however, perfect, and the quality of variants detected by sequencing may be adversely influenced by several factors. First, genome sequencing is known to have errors or biases(17–21), which might cause inaccuracy in detecting variants. Second, sequence mappability of different regions may not be uniform, but correlated with sequence-specific biological features, leading to alignment biases. For instance, it is shown that introns have significantly lower mappability levels than exons(22). Third, variant calling algorithms may be sources of errors as no algorithm is 100% accurate. For example, GATK HaplotypeCaller and GATKUnifiedGenotyper(23), which are the widely used variant callers, have sensitivity of about 96% and precision of about 98%(24). Additionally, different variant callers may generate discordant calls on some variants(25), which indicates inaccuracy of those calls, and in certain cases, different versions of even the same software may generate inconsistent calls. All these factors may generate false positive sites or incorrect genotypes, which may then lead to false positive associations in the follow-up association test. For example, Alzheimer’s Disease Sequencing Project reports that they found spurious associations in the case-control analysis where one of the causes for the problem could be inconsistent variant calling processes for sequenced samples(26).

It is extremely important to perform quality control (QC) on genetic variants identified from sequencing to remove variants that may contain sequencing errors and hence are likely to be false positive calls. Traditionally, genetic studies have utilized two types of QC approaches; we call them, “filtering” and “classification” approaches. In filtering approaches, several filters are applied to remove problematic variants such as variants with high genotype missing rate (e.g. > 5%), low Hardy-Weinberg Equilibrium (HWE) p-value (e.g. < 1E-4), or very high or low allele balance of heterozygous calls (ABHet) (e.g. > 0.75 or < 0.25). One main problem with this type of approaches is that these thresholds are arbitrarily determined without strong statistical justification. We may also remove variants whose metrics are very close to the thresholds (e.g. variants with missing rate of 5.1%). Another type of QC is a classification approach that attempts to learn variants with low quality using machine learning approaches. One example is VQSR of GATK(24, 27) that uses a Gaussian mixture model to learn the multidimensional annotation profile of variants with high and low quality. However, one of issues with VQSR is that one needs training datasets acquired from existing databases on variants such as 1000 Genomes Project(28) and HapMap(29), which may be biased to keep known variants and filter out novel variants. Another issue is that those known databases of genetic variants may not be always accurate, which would lead to inaccurate classification of variants, and they may not even be available for some species. It may also be a challenge to apply VQSR to a variant call set generated by variant callers other than GATK as VQSR needs metrics of variants that are not often calculated by non-GATK variant callers.

In this article, we present ForestQC for performing QC on genetic variants discovered through sequencing. Our method aims to identify whether a specific variant is of high sequencing quality (“good” variants) or of low quality (“bad” variants) by combining the filtering and classification approaches. We first apply a filtering approach to detect obviously good and bad variants from data. We use stringent filters such that those variants are truly good or bad while the rest of variants that are neither good nor bad are considered to have ambiguous quality (“gray” variants). Given this set of good and bad variants, we train a machine learning model whose goal is to classify whether gray variants are good or bad. With an insight that good variants would have higher genotype quality and sequencing depth than do bad variants, we use information of several sequencing quality measures of variants for model training. ForestQC then uses sequencing quality measures of gray variants to predict whether each gray variant has high or low sequencing quality. Our approach is different from the filtering strategy in that it only uses filters to identify truly good or bad variants and does not attempt to classify gray variants with filters. Our method is also different from VQSR as our training strategy allows us to train our model without known datasets for variants and solves several issues with VQSR mentioned above. Another advantage of our software is that it can be applied to standard Variant Call Format (VCF) files from any variant callers and is very efficient.

To demonstrate accuracy of ForestQC, we apply it to two high-coverage WGS datasets; 1) large extended pedigrees ascertained for bipolar disorder (BP) from Costa Rica and Colombia(30), and 2) a sequencing study for Progressive Supranuclear Palsy (PSP). The first dataset includes 449 related individuals from families while the latter dataset consists of 495 unrelated individuals. We show that ForestQC outperforms VQSR and a filtering approach based on ABHet as good variants detected from ForestQC have higher sequencing quality than those from VQSR and the filtering approach in both datasets. This suggests that our tool identifies high-quality variants more accurately than other approaches in both family and unrelated datasets. ForestQC is publicly available at https://github.com/avallonking/ForestQC

## Results

### Overview of ForestQC

ForestQC takes a raw VCF file as input and determines whether each variant has “good” sequencing quality or “bad” quality. Our method combines a filtering approach that determines good and bad variants by a set of pre-defined filters and a classification approach that uses machine learning to classify whether a variant is good or bad. As illustrated in Figure 1, our method first calculates statistics of each variant for several filters that are commonly used in performing QC in GWAS. These statistics consist of ABHet, HWE p-value, genotype missing rate, Mendelian error rate for family data, and any user-defined statistics (details described in Method session). ForestQC then identifies three sets of variants using these statistics for filters: 1) a set of good variants that pass all filters, 2) a set of bad variants that fail any filter(s), and 3) a set of gray variants that are neither good nor bad variants. We use stringent thresholds for filters (Table S2, S3), and hence we are highly confident that good variants are of high quality while bad variants are truly false positives or have unequivocally poor sequencing quality. The next step in ForestQC is to train a random forest machine learning model using the good and bad variants we detect from the filtering step. In ForestQC, seven sequencing quality metrics of good and bad variants are used as features to train the random forest model, including three related to sequencing depth, three related to genotype quality, and one related to the GC content. Finally, the fitted model predicts whether each gray variant is good or bad. We combine the predicted good variants from the random forest model and the good variants from the filtering step, and they are all good variants determined by ForestQC. The same procedure is applied to identify bad variants.

**Figure 1:**
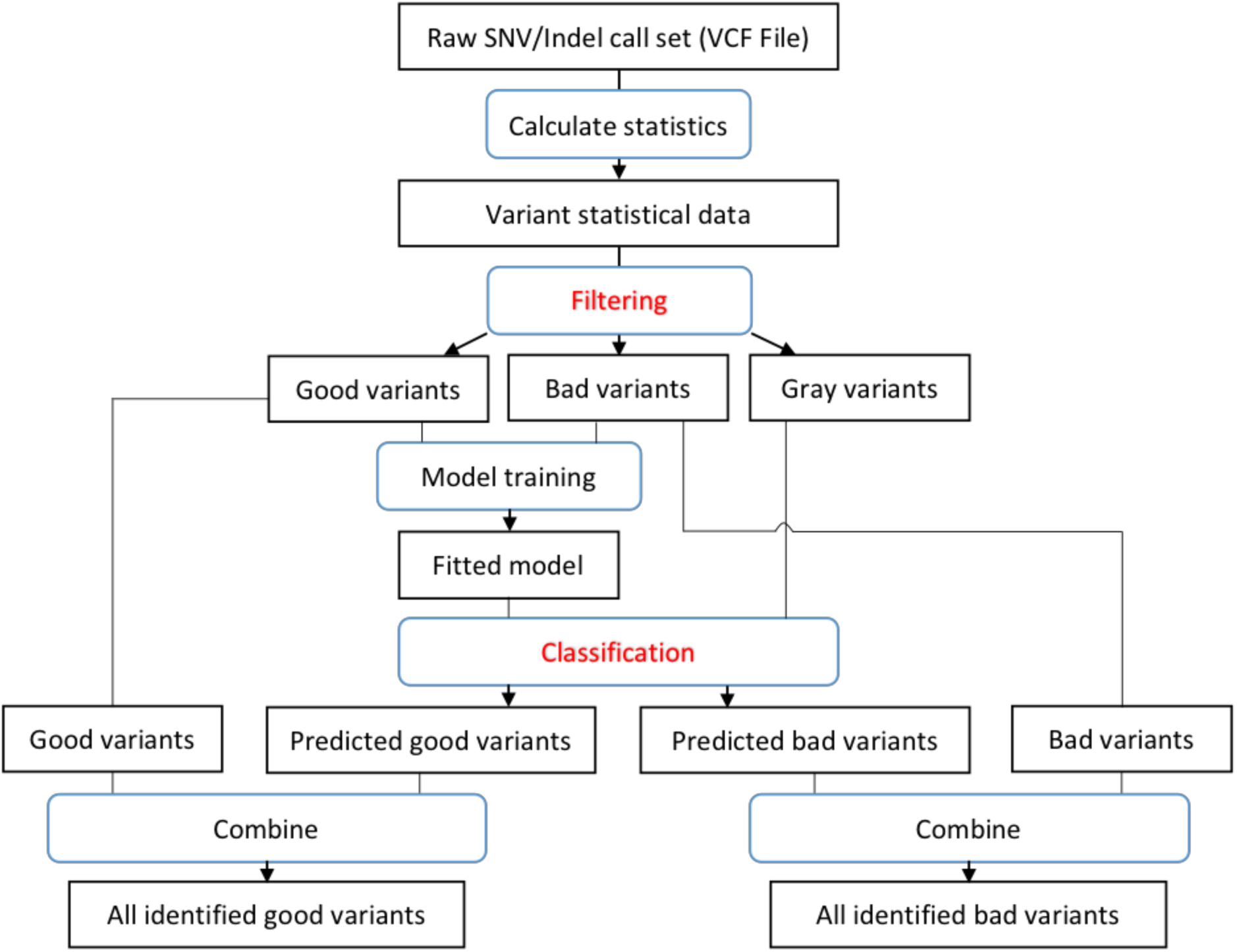
Workflow of ForestQC. ForestQC takes a raw variant call set in the VCF format as input. Then it calculates the statistics of each variants, including MAF, mean depth, mean genotyping quality, etc.. In the filtering step, it separates the variant call set into good, bad, and gray variants by applying various hard filters, such as Mendelian error rate and genotype missing rate. In classification step, good and bad variants are used to train a random forest model, which is then applied to assign labels to gray variants. Variants predicted to be good among gray variants are combined with good variants from the classification step for the final set of good variants. The same procedure applies to find the final set of bad variants.

One major challenge in classifying gray variants is to identify a set of sequencing quality metrics that are used as features to train the random forest model. We choose three sets of features based on quality metrics that variant callers provide and prior knowledge in genome sequencing. The first set of features is genotype quality (GQ) where we have three metrics: mean, standard deviation (SD), and outlier ratio. The outlier ratio is the proportion of samples whose GQ scores are lower than a particular threshold, and it measures a fraction of individuals who are poorly sequenced at a mutation site. A good variant is likely to have high mean, low SD, and low outlier ratio of GQ values. The second set of features is sequencing depth (DP) as low depth often introduces sequencing biases and reduces variant calling sensitivity(31). We also use the same three sets of metrics for DP as those for GQ: mean, SD, and outlier ratio. The last set of features is related to genomic characteristics instead of sequencing quality, which is GC content. High or low GC content may decrease the coverage of certain regions(32, 33) and thus may lower the quality of variant calling. Hence, the GC content of the DNA region containing a good variant would not be too high or too low. Given these three sets of features, ForestQC learns how those features determine good and bad variants and classifies gray variants according to rules that it learns.

### Comparison of different machine learning algorithms

As there are many different machine learning algorithms available, we first seek to find the most accurate and efficient algorithm for performing QC on NGS variant data. To ensure the quality of training and prediction, we choose supervised learning algorithms rather than unsupervised algorithms. Several major types of supervised algorithms are selected for comparison: random forest, logistic regression, k nearest neighbors (KNN), Naive Bayes, quadratic discriminant analysis (QDA), AdaBoost, artificial neural network (ANN), and single support vector machine (SVM). We use the BP WGS dataset, which consists of large pedigrees from Costa Rica and Colombia, to compare the performance of different algorithms. We use the aforementioned three sets of features related to sequencing quality for all algorithms we test. We apply the filtering approach (Table S2, S3) to the BP data to identify good, bad, and gray variants, and we choose 100,000 good and 100,000 bad variants randomly for model training. We then choose another 100,000 good and 100,000 bad variants randomly from the rest of variants for model testing. Each learning algorithm will be trained with the same training set and tested with the same test set. We use 10-fold cross validation, area under the receiver operating characteristic curve (AUC), and F1-score to estimate classification accuracy during model testing. F1-score is the harmonic average of precision (positive predictive value) and recall (sensitivity). The closer F1-score is to 1, the better the performance is. To assess the efficiency of each algorithm, we measure its time cost during training and predicting. We use eight threads for algorithms that support parallelization.

Results show that random forest is the most accurate model in both SNV classification and indel classification with the highest F1-scores, accuracy and the largest AUC (Table 1, Table S1, Figure S1). Its time cost is only 9.85 seconds in model training and prediction (Table 1), which ranks as the fourth fastest algorithm. As random forest randomly divides the entire dataset into several subsets of the same size and constructs decision trees independently in each subset, it is highly scalable, and it has low error rates and high robustness with respect to noise(34). As for other machine learning algorithms, both SVM and ANN are highly accurate (both with F1-score of 0.97 and AUC > 0.985 in SNV classification) but they are not as efficient as random forest. ANN is the second slowest algorithm that is about 8x slower than random forest because it has to estimate many parameters. Especially, SVM is the slowest algorithm because of its inability to parallelize, which costs about 125x as much time as random forest (Table 1). This suggests that it may be computationally very expensive to use SVM in large-scale WGS datasets that have tens of millions of variants. Normally, a real dataset is at least 10 times larger than the dataset used here. For example, in the BP dataset, the training set has 2.20 million (M) SNVs and there are 2.73M gray SNVs for prediction. We find that random forest only spends 80.51 seconds for training and predicting, while ANN needs 489.63 seconds and SVM needs 14.74 hours. Therefore, random forest is much faster than ANN and SVM, although all three algorithms have similar performance in terms of AUC (Figure S1). In addition, there are even a larger number of variants in large-scale WGS projects such as NHLBI Trans-Omics for Precision Medicine (TOPMed) program that includes about 463M variants. Hence, it is more practical to use random forest when processing this very large datasets. Logistic regression, Naive Bayes and QDA are more efficient than random forest, but their predictions are not as accurate as those of random forest. For example, Naive Bayes needs only 0.18 seconds for training and prediction while its F1-score is the lowest among all algorithms (0.90 and 0.87 in SNV and indel classification, respectively) (Table 1). This result demonstrates that random forest is both accurate and efficient, and hence we use it as the machine learning algorithm in our approach. To further improve the random forest algorithm, we test a different number of trees in the algorithm and we find that random forest with 50 trees balances efficiency and accuracy (Figure S2). To identify good variants from gray variants, we use the probability of each gray variant being a good variant calculated from random forest, and we consider gray variants with the probability of being good variants > 50% as good variants as this probability threshold achieves the highest F1-score (Figure S3).

**Table 1:**
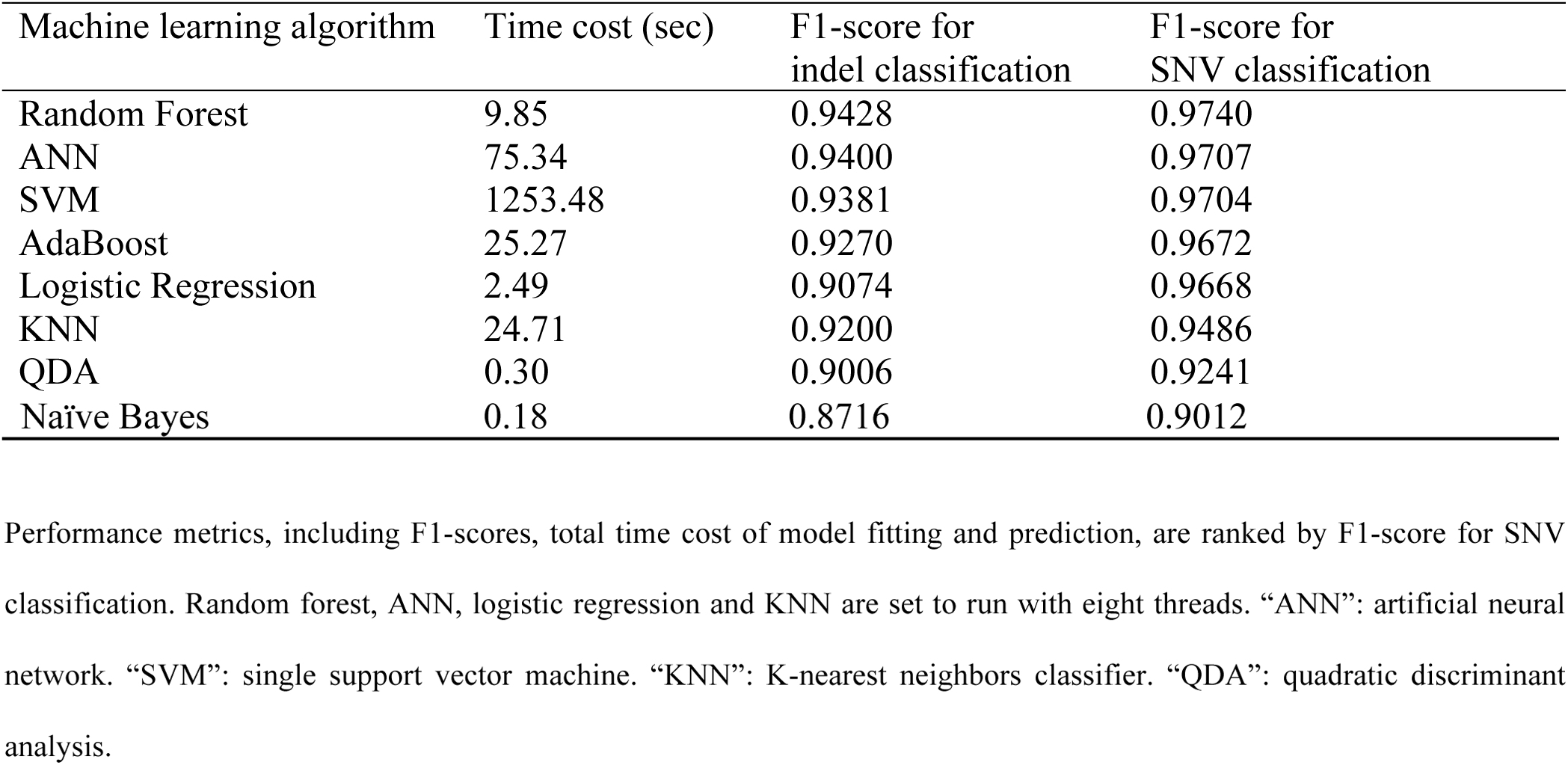
Performance of eight different machine learning algorithms

### Measuring performance of QC methods on WGS data

To evaluate the accuracy of ForestQC and other methods on WGS data, we apply them to two WGS datasets and calculate several statistics. For a family-based dataset, we calculate Mendelian error rate (ME) of each variant, which measures inconsistency in genotypes between parents and offspring. Another statistic we measure is genotype discordance rate between microarray and sequencing if individuals who are sequenced are also genotyped. In both WGS datasets we analyze, microarray data are available. These two statistics are important indicators of quality of variants because good variants would follow Mendelian inheritance patterns and their genotypes would be consistent between microarray and sequencing. In addition to these statistics, we measure several other statistics that are reported in sequencing studies such as the number of variants (SNVs and indels), transitions/transversions (Ti/Tv) ratio, the number of multi-allelic variants, genotype missing rate. We compute these QC-related statistics separately for SNVs and indels. We use these statistics to compare the performance of ForestQC with that of three approaches. The first is one without performing any QC (no QC). The second method is VQSR which is a classification approach that requires known truth sets for model training, such as HapMap or 1000 genomes. We use recommended resources and parameter settings to run VQSR as of 2018-04-04(35), but we also look at different settings. The third method is an ABHet approach, which is a filtering approach that retains variants according to allele balance of variants (see Methods).

### Performance of ForestQC on family WGS data

We apply ForestQC to the BP WGS dataset that consists of 449 subjects with the average coverage of 36. There are 25.08M SNVs and 3.98M indels(30). The variant calling is performed with GATK-HaplotypeCaller v3.5. This is an ideal dataset for assessing the performance of different QC methods because this dataset contains individuals from families who are both sequenced and genotyped. This study design allows us to calculate both ME rate and genotype discordance rate of variants between WGS and microarray. For this dataset, we test ForestQC with two different filter settings, one using ME rate as a filter and the other not using ME as a filter. The results of the former approach would filter out bad variants based on ME rate, and hence ME rate of good variants would be very low. However, we observe that both approaches have similar performance in terms of ME rate and other statistics (Table S4, Figure S4, Figure S5), and hence we show results of only ForestQC using ME rate as a filter.

Results show that ForestQC outperforms ABHet and VQSR in terms of the quality of good SNVs while it detects fewer good SNVs than the other approaches (detailed variant-level metrics in Table S5). ForestQC identifies 22.23M (88%) good SNVs, which is fewer than 22.42M (89%) and 24.24M (97%) good SNVs from ABHet and VQSR, respectively (Table 2). However, ABHet has 3.57x and VQSR has 9.99x higher ME rate on good SNVs than ForestQC (Figure 2a), and ABHet has 1.50x (p-value < 2.2e-16) and VQSR has 1.26x higher genotype discordance rate (p-value < 2.2e-16) on good SNVs than ForestQC (Figure 2b). In addition, ABHet and VQSR have 81.48x and 97.72x higher genotype missing rate on good SNVs than ForestQC, respectively (Figure 2c), but it is important to note that genotype missing rate is used as a filter in ForestQC, which means SNVs with high genotype missing rate are filtered out. We observe that VQSR and ABHet have 319 thousand (K) (1.32%) and 235K (1.05%) good SNVs with very high genotype missing rate (>10%), respectively, and there are also 118K (0.49%, VQSR) and 53K (0.24%, ABHet) good SNVs with very high ME rate (>15%) while ForestQC has none of them due to its filtering approach. The better quality of good SNVs from ForestQC means that bad SNVs detected from ForestQC would have lower quality, and results show that bad SNVs detected by our method have higher genotype missing rate, higher ME rates and higher genotype discordance rate than those of ABHet, and higher genotype missing rate than those of VQSR (Figure S6a, b, c). The no QC method keeps the greatest number of good SNVs (25.08M), but they have the highest ME rate, genotype missing rate, and genotype discordance rate as expected.

**Figure 2:**
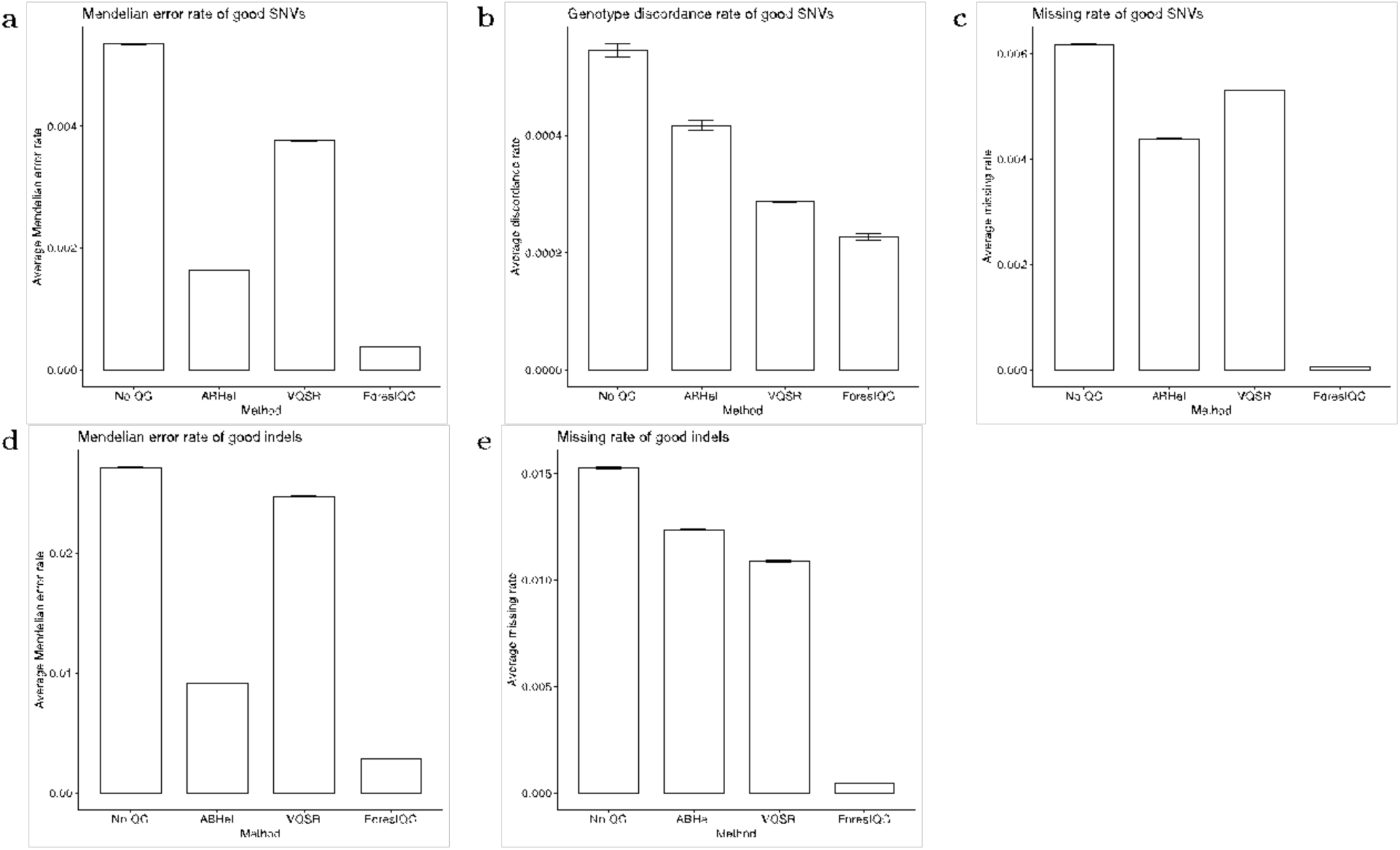
Overall quality of good variants in the BP dataset detected by four different methods, including no QC applied, ABHet approach, VQSR and ForestQC. The average Mendelian error rate and genotype missing rate for SNVs and indels, and genotype discordance rate to microarray data for SNVs are shown. Data are represented as the mean ± SEM.

**Table 2:**
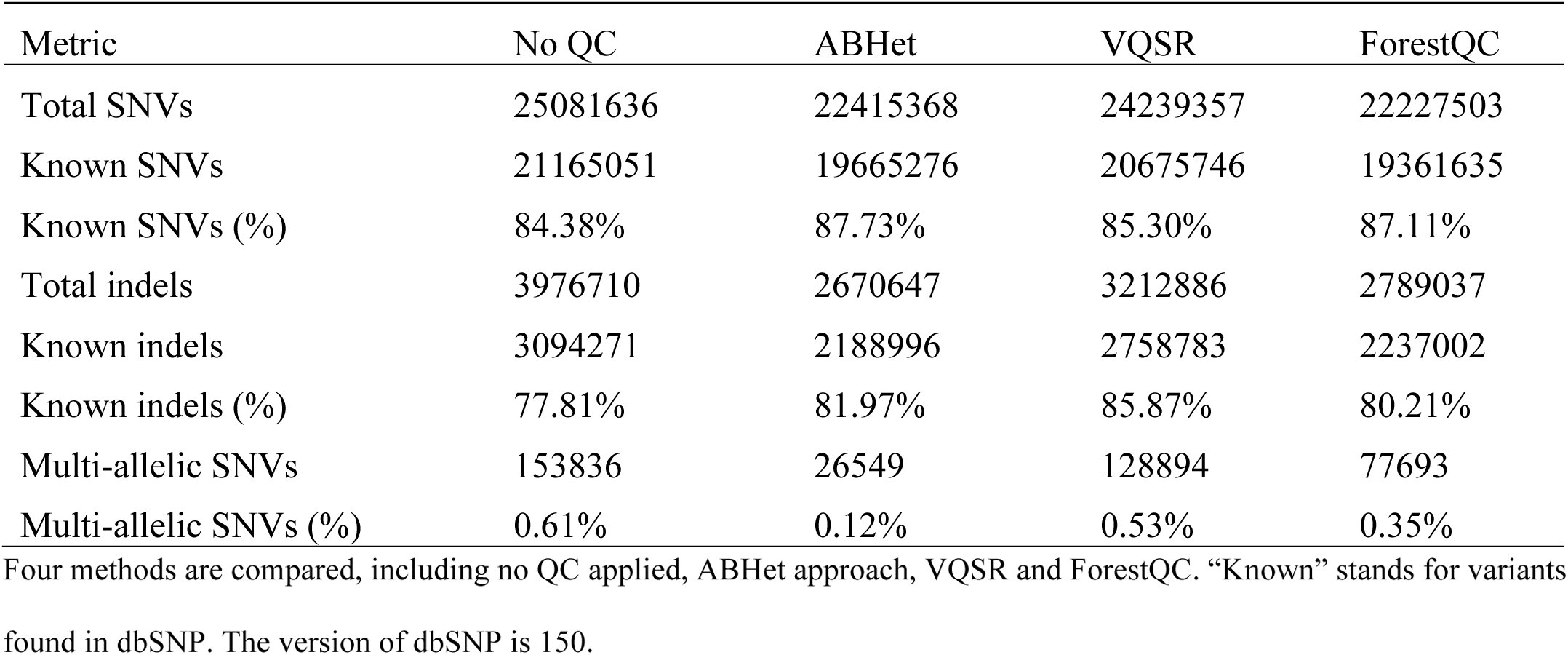
Variant-level quality metrics of good variants in the BP dataset processed by different methods

Next, we obtain several statistics of good SNVs commonly used in sequencing studies to evaluate the performance of ForestQC. One such statistic is Ti/Tv ratio, which is expected to be around 2.0 over the whole genome(36). If this ratio is smaller than 2.0, it means that there may be false positive variants in the dataset. We compute Ti/Tv ratio for each individual across all good SNVs and look at the distribution of those ratios across all individuals (sample-level statistics). We find that the mean Ti/Tv ratio of good known SNVs (present in dbSNP) is around 2.0 for all four methods, which suggests that they have similar accuracy on known SNVs in terms of Ti/Tv ratio (Figure S7a). However, results show that the mean Ti/Tv ratio of good novel SNVs (not in dbSNP) from ForestQC is better than that of those SNVs from other methods; the mean Ti/Tv ratio is 1.68 for ForestQC, which is closest to 2.0 among other methods (1.41 for VQSR, 1.53 for ABHet, and 1.29 for No QC) (Figure 3a). Paired t-tests for the difference in the mean Ti/Tv ratio between ForestQC and other methods are all significant (p-value < 2.2e-16 versus all other methods). This result suggests that novel SNVs predicted to be good by ForestQC are more likely to be true positives than those SNVs from other QC methods. Another statistic commonly used in sequencing studies is the percentage of multi-allelic SNVs, which are variants with more than one alternative allele. Given this sample size (449), many of them are likely to be false positives, and ForestQC has 33.96% and 42.62% smaller fraction of multi-allelic SNVs among good SNVs than do VQSR and no QC methods while the ABHet approach has the smallest fraction of such SNVs (Table 2). Note that ABHet values can only calculated for biallelic mutation sites, so ABHet does not work properly for multi-allelic variants. It might mistakenly filter out many high quality multi-allelic SNVs, so it has the fewest multi-allelic SNVs.

**Figure 3:**
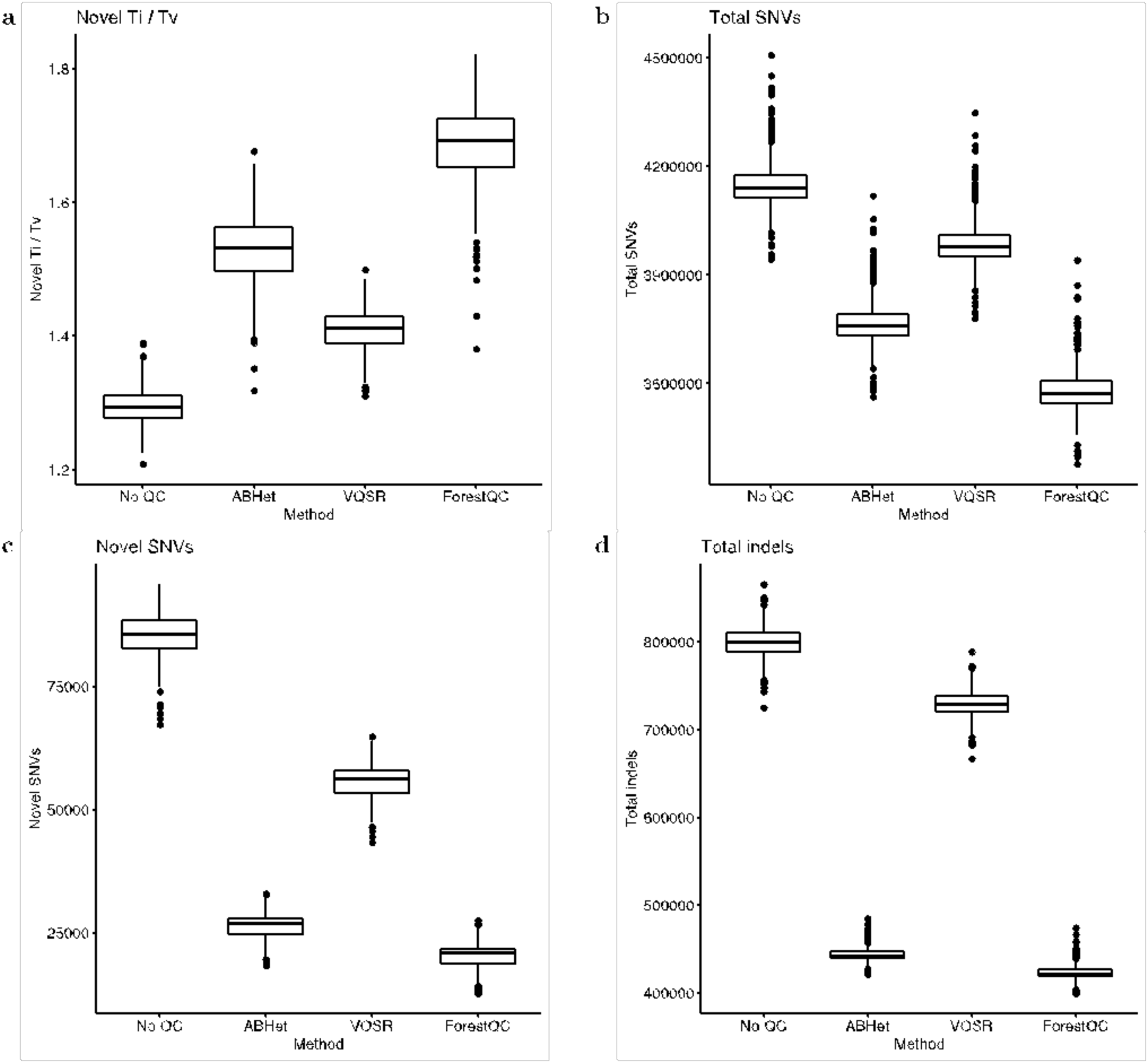
Sample-level quality metrics of good variants in the BP dataset identified by four different methods, including no QC applied, ABHet approach, VQSR and ForestQC. (a) Ti/Tv ratio of SNVs not found in dbSNP. (b) Total number of SNVs. (c) The number of SNVs not found in dbSNP. (d) Total number of indels. The version of dbSNP is 150.

In addition to SNVs, we apply the four QC methods to indels. Similar to results of SNVs, ForestQC identifies fewer good indels than does VQSR, but the quality of those indels from ForestQC is better than that of good indels from ABHet and VQSR. Out of total 3.98M indels, ForestQC predicts 2.79M indels (70%) to have good sequencing quality while VQSR and ABHet find 3.21M (81%) and 2.67M (67%) good indels, respectively (Table 2). Good indels from VQSR and ABHet, however, have 8.54x and 3.18x higher ME rate, and 22.25x and 25.28x higher genotype missing rate, than those from ForestQC, respectively (Figure 2d, e). Bad indels identified by ForestQC have 2.25x and 1.32x higher ME rate, and 1.48x and 2.36x higher genotype missing rate than those from VQSR and ABHet, respectively (Figure S6d, e). Besides, we observe that there are 95K (2.97%, VQSR) and 86K (3.23%, ABHet) good indels with very high genotype missing rate (>10%) and also 167K (5.21%, VQSR) and 44K (1.66%, ABHet) good indels with very high ME rate (>15%) while there are no such indels in ForestQC. This result suggests that many good indels detected by ABHet or VQSR may be false positives or indels with poor sequencing quality. One of the reasons why VQSR does not perform well on indels could be the database it uses for training its machine learning model as VQSR considers all indels found in the database (Mills gold standard call set(37) and 1000G Project(38)) to be true variants. This leads VQSR to have a significantly higher proportion of known indels among good indels (86%), compared with 80% from ForestQC and 82% from ABHet (Table 2). The poor performance of VQSR on indels may be because not all indels in the database are true variants, or because even if they are true indels, those indels would not necessarily have high sequencing quality in the sequencing dataset of interest. Hence, this result demonstrates one of the limitations of using known databases for finding good variants. It is also important to note that in general, indels have much higher ME rate (0.41% for no QC) than that of SNVs (0.08% for no QC), which is expected given the greater difficulty of calling indels.

Another major difference between ForestQC and the other approaches is the allele frequency of variants after QC as ForestQC keeps a greater number of rare variants in its good variant set. Our method has 1.77% and 1.64% higher proportion of rare SNVs, and 5.30% and 15.37% higher proportion of rare indels than ABHet and VQSR do, respectively (Table S6). We also observe this phenomenon in the variant-level and sample-level statistics for the number of SNVs. The variant-level statistics show that the number of good SNVs detected by ForestQC is similar to those from ABHet (Table 2). However, the sample-level statistics show that each individual on average carries fewer alternative alleles of good SNVs from ForestQC (3.58M total SNVs) than those from VQSR and ABHet (3.99M and 3.77M total SNVs, respectively) (Figure 3b, c, Figure S7b). We observe a similar phenomenon for indels between ABHet and ForestQC (Table 2, Figure 3d, Figure S7c, d). This phenomenon could be explained by the higher fraction of rare variants among good variants from ForestQC, as individuals would carry fewer variants if there are a greater fraction of rare variants. One main reason why ForestQC has the higher proportion of rare variants is that common variants have higher ME rate, genotype discordance rate and genotype missing rate than do rare variants (Figure S8); because common variants are more heterozygous, it is more difficult to accurately call them. This suggests that while a majority of common variants may be true variants, some of them may not necessarily have high sequencing quality, and hence their calls may not be accurate enough for downstream analyses.

ForestQC uses several filters to remove variants whose sequencing quality is poor while other two approaches (VQSR and ABHet) do not use these filters, which might have artificially improved the performance of ForestQC. Hence, to compare the performance of ForestQC with other approaches without this potential bias due to the filtering step, we measure the performance metrics on only gray variants as their sequencing quality is not determined by the filtering approach. From 2.73M gray SNVs and 1.09M gray indels, ForestQC identifies 979K (35.83%) good SNVs and 532K (48.58%) good indels, while ABHet approach detects 620K (22.70%) SNVs and 195K (17.80%) indels, and VQSR selects 2.16M (79.18%) SNVs and 643K (58.76%) indels as good variants, respectively (Table S7). For good SNVs from gray variants, ABHet and VQST have 2.75x and 22.67x higher ME rate than ForestQC, respectively (Figure S9a), and ABHet and VQSR have 5.15x (p-value = 1.367e-14) and 3.86x (p-value = 1.926e-14) higher genotype discordance rate than ForestQC (Figure S9b). In addition, ABHet and VQSR have 15.50x and 7.05x higher genotype missing rate on good SNVs than ForestQC, respectively (Figure S9c). We observed similar results for indels (Figure S9d and S8e). Sample-level metrics also show that ForestQC has better Ti/Tv ratio on known SNVs (mean Ti/Tv: 1.64, 1.85, 1.72, 1.88 for No QC, ABHet, VQSR, ForestQC, respectively), and novel SNVs (mean Ti/Tv: 1.14, 1.04, 1.21, 1.22 for No QC, ABHet, VQSR, ForestQC, respectively) than other methods (Figure S10d and S9e). Paired t-tests for the difference in the mean Ti/Tv ratio of novel SNVs and known SNVs between ForestQC and other methods are all significant (p-value < 0.05 versus all other methods). These results show that even on those variants for whom we do not use the filtering approach, ForestQC has better performance than ABHet and VQSR. These results further imply that if we use the same filtering approach to all three approaches, our method will still outperform other approaches.

### Performance of ForestQC on WGS data with unrelated individuals

To evaluate the performance of ForestQC on WGS datasets that contain only unrelated individuals, we apply it to the PSP dataset that has 495 individuals who are whole-genome sequenced at average coverage of 29, generating 33.27M SNVs and 5.09M indels. Among the 495 individuals who are sequenced, 381 individuals (77%) of them are also genotyped with microarray, which enables us to check the genotype discordance rate between WGS and microarray data. Because the PSP dataset contains only unrelated individuals, we do not report ME rate. Similar to BP WGS data, we apply four methods (ForestQC, VQSR, ABHet, and No QC) to the PSP dataset, although the parameter setting of VQSR has slightly changed. As the PSP dataset is called with GATK v3.2, the StrandOddsRatio (SOR) information from the VCF file is missing, which is recommended to use in VQSR, and hence this annotation is excluded from VQSR. However, we find that SOR information has little impact on the results of VQSR as we test VQSR without SOR information using the BP dataset and obtain similar results with one using SOR information (Figure S11).

Similar to the results of the BP dataset, ForestQC identifies good variants with higher quality although it detects fewer good variants than other approaches (detailed variant-level metrics in Table S8). ForestQC identifies 29.25M (88%) good SNVs, which is slightly fewer than 29.77M (89%) good SNVs from ABHet but about 2 million fewer than 31.28M (94%) good SNVs from VQSR (Table 3). However, good SNVs from ABHet and VQSR have 53.76x and 42.55x higher genotype missing rate than those from ForestQC, respectively (Figure 4a), but it is important to note that missing rate is included as a filter in ForestQC. In addition, there are 311K (0.99%, VQSR) and 331K (1.13%, ABHet) good SNVs with very high genotype missing rate (>10%), while ForestQC removes all these SNVs. We also observe that bad SNVs from ForestQC have 2.4x higher genotype missing rate than those from ABHet, although bad SNVs from GATK have slightly higher missing rate than those from ForestQC (Figure S12a). Good SNVs from ABHet and VQSR have 1.28x (p-value < 2.2e-16) and 1.29x higher genotype discordance rate (p-value < 2.2e-16) than those from ForestQC, respectively (Figure 4b). As for the genotype discordance rate of bad SNVs, both ABHet and VQSR have higher genotype discordance rate than does ForestQC (Figure S12b), but this may be inaccurate because of the small number of bad SNVs genotyped with microarray (10,130, 4,121, and 553 such SNVs for ForestQC, ABHet, and VQSR, respectively). The variant-level and sample-level statistics also demonstrate the better quality of good SNVs from ForestQC. Although all methods have mean Ti/Tv ratio of good known SNVs above 2.0, the mean Ti/Tv ratio of good novel SNVs among all sequenced individuals is 1.65 for ForestQC, which is closer to 2.0 than other methods (1.27, 1.54, and 1.24 for VQSR, ABHet, no QC, respectively). (Figure S13a, Figure 5a). Paired t-tests for the difference in the mean Ti/Tv ratio between ForestQC and other methods are all significant (p-value < 2.2e-16 versus all other methods). ForestQC has 16.67% and 33.33% smaller fraction of multi-allelic SNVs among good SNVs than do VQSR and no QC methods, respectively, while the ABHet approach has the smallest proportion of such SNVs (Table 3). ABHet has the smallest number of multi-allelic SNVs because it can only work properly for biallelic SNVs where all subjects are either heterozygous or homozygous and therefore it might remove many multi-allelic SNVs by mistakes. Lastly, consistent with the results of the BP dataset, the sample-level statistics show that each individual on average carries fewer alternative alleles of good SNVs from ForestQC than those from VQSR and ABHet (Figure 5b, c, Figure S13b). Rare SNVs in good SNVs from ForestQC account for 1.70% and 1.32% higher proportion, compared with those from ABHet and VQSR (Supplemental Table 5). This may be because rare SNVs have lower genotype missing rate and genotype discordance rate than do common variants (Figure S14a, b).

**Figure 4:**
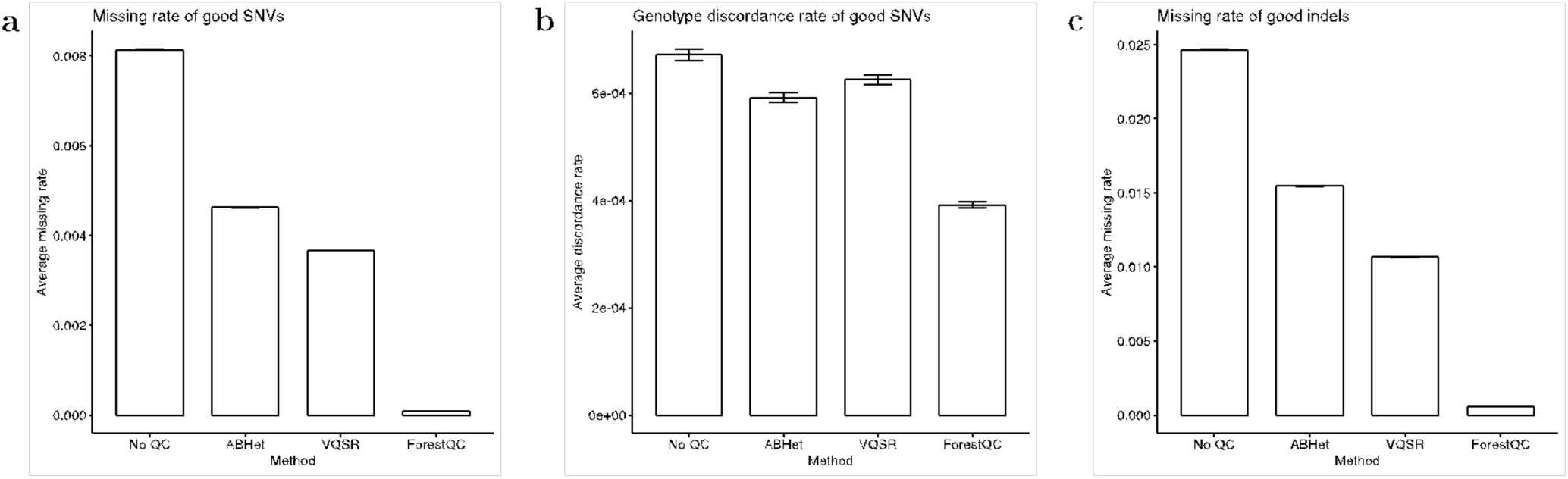
Overall quality of good variants in the PSP dataset detected by four different methods, including no QC applied, ABHet approach, VQSR and ForestQC. The average genotype missing rate for both SNVs and indels, and genotype discordance rate to microarray data for SNVs are shown. Data are represented as the mean ± SEM.

**Figure 5:**
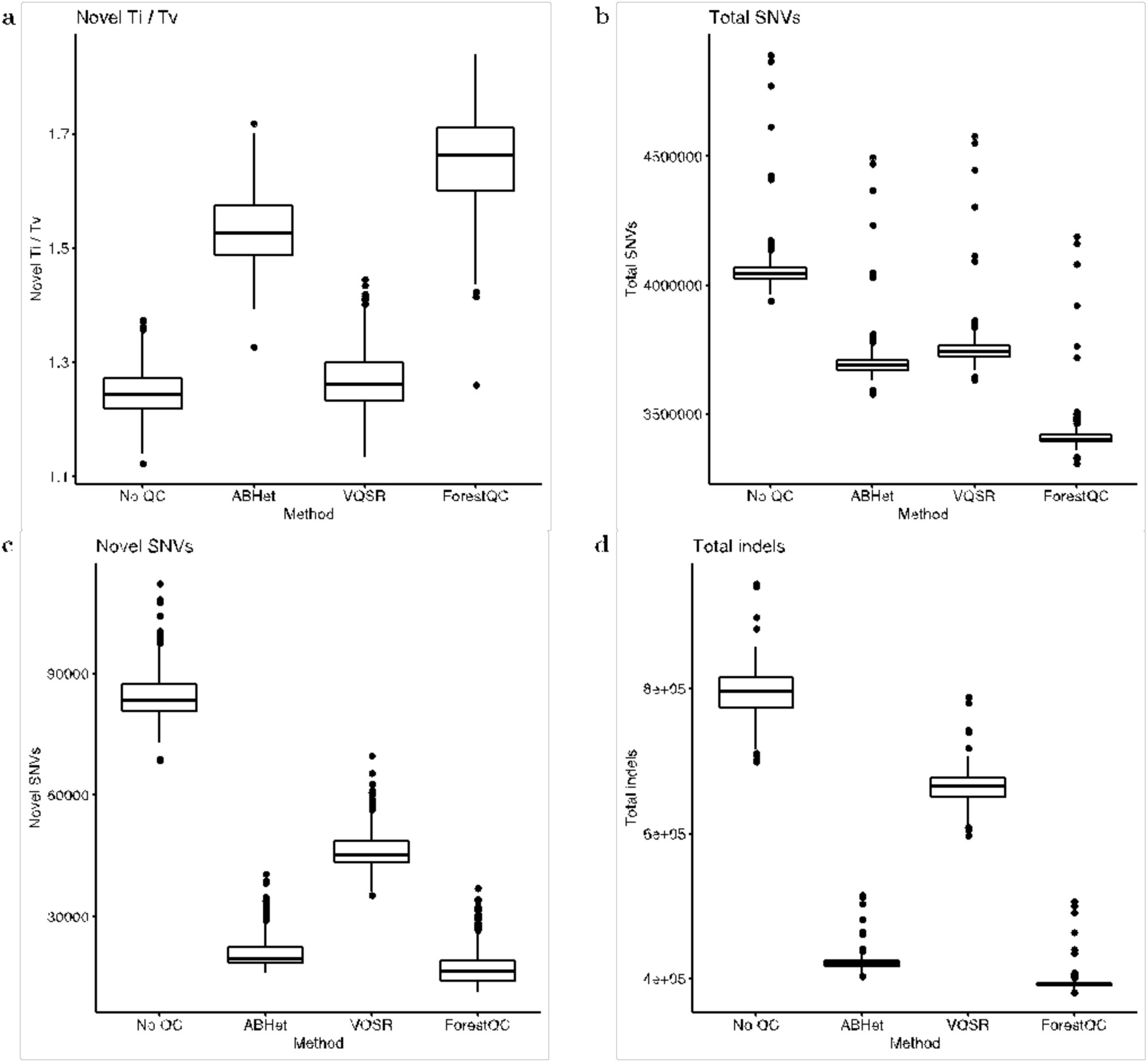
Sample-level quality metrics of good variants in the PSP dataset identified by four different methods, including no QC applied, ABHet approach, VQSR and ForestQC. (a) Ti/Tv ratio of SNVs not found in dbSNP. (b) Total number of SNVs. (c) The number of SNVs not found in dbSNP. (d) Total number of indels. The version of dbSNP is 150.

**Table 3:**
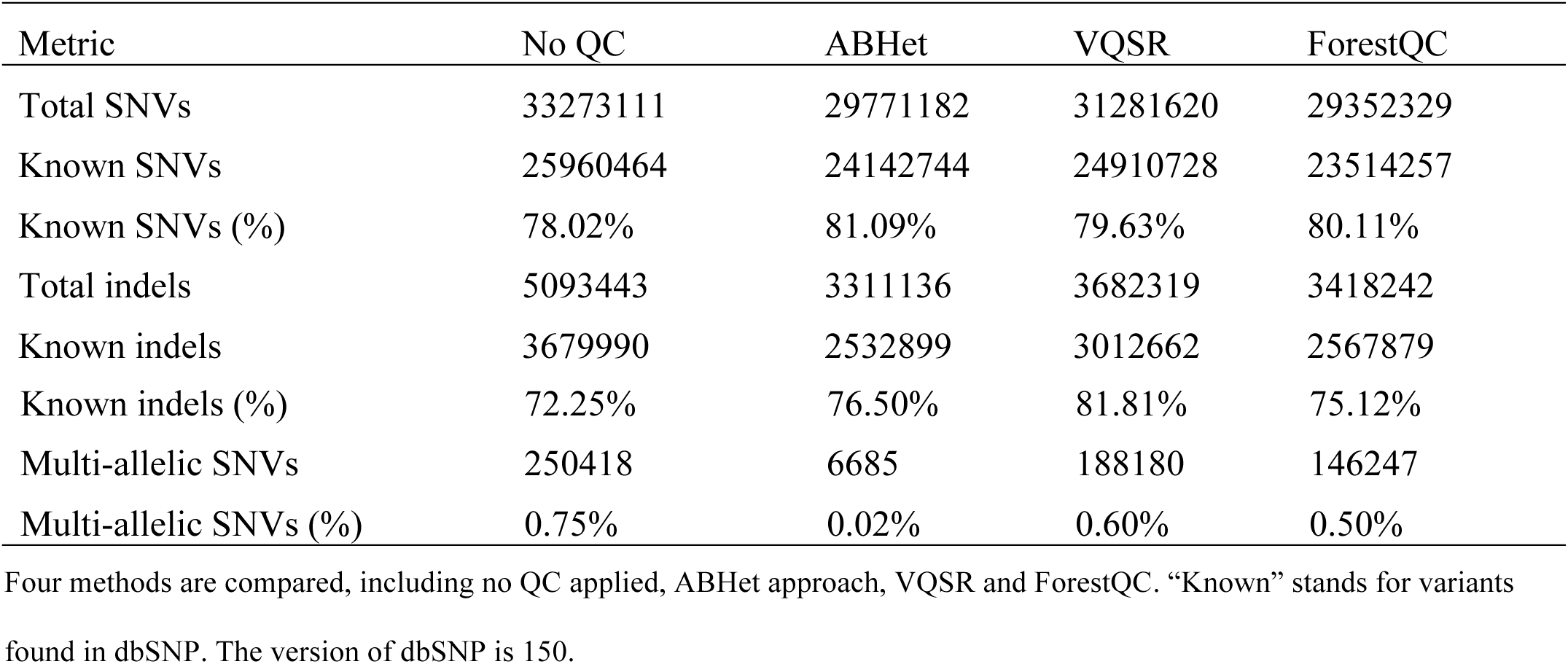
Variant-level quality metrics of good variants in the PSP dataset processed by four different methods

| Metric | No QC | ABHet | VQSR | ForestQC |
| --- | --- | --- | --- | --- |
| Total SNVs | 33273111 | 29771182 | 31281620 | 29352329 |
| Known SNVs | 25960464 | 24142744 | 24910728 | 23514257 |
| Known SNVs (%) | 78.02% | 81.09% | 79.63% | 80.11% |
| Total indels | 5093443 | 3311136 | 3682319 | 3418242 |
| Known indels | 3679990 | 2532899 | 3012662 | 2567879 |
| Known indels (%) | 72.25% | 76.50% | 81.81% | 75.12% |
| Multi-allelic SNVs | 250418 | 6685 | 188180 | 146247 |
| Multi-allelic SNVs (%) | 0.75% | 0.02% | 0.60% | 0.50% |

Four methods are compared, including no QC applied, ABHet approach, VQSR and ForestQC. “Known” stands for variants found in dbSNP. The version of dbSNP is 150.

For indels, our method predicts 3.42M indels (67% of total 5.09M indels) to be good variants, which is slightly more than 3.31M (65%) good indels from ABHet and fewer than 3.68M (72%) good indels from VQSR (Table 3). Because the PSP dataset lacks ME rate as it contains only unrelated individuals and indels are not called in microarray, it is difficult to compare the performance of the QC methods on indels. We find that good indels from ABHet and VQSR have 27.02x and 18.77x higher genotype missing rate than those from our method, respectively (Figure 4c). Additionally, VQSR and ABHet have 107K (2.91%) and 131K (4.08%) good indels with high genotype missing rate (>10%), respectively while ForestQC filters out all of these indels. Also, bad indels from ForestQC have 2.05x and 1.21x higher genotype missing rate than those from ABHet and VQSR, respectively (Figure S12c). This, however, may be biased comparison as ForestQC removes indels with high genotype missing rate in its filtering step. Consistent with the results of SNVs, the sample-level statistics indicate that each individual has fewer good indels from ForestQC than those from VQSR and ABHet (Figure 5d, Figure S13c, d). Among good indels, ForestQC has 6% and 1% more novel indels than VQSR and ABHet, respectively (Table 3). In terms of allele frequency, rare indels detected by ForestQC accounts for 12.35% and 3.49% larger proportions than those by VQSR and ABHet, respectively (Table S9). Similar to the results of the BP dataset, we also observe that the missing rate of rare indels is lower than that of common indels (Figure S14c).

Similar with the analysis of the BP dataset, we also compare the performance of ForestQC, ABHet approach and VQSR only on gray variants in PSP dataset. From 3.95M gray SNVs and 1.60M gray indels, ForestQC identifies 1.71M (43.33%) good SNVs and 719K (45.01%) good indels, while ABHet approach detects 780K (19.74%) SNVs and 248K (15.51%) indels, and VQSR selects 2.75M (69.52%) SNVs and 820K (51.34%) indels as good variants, respectively (Table S10). For good SNVs from gray variants, ABHet and VQSR have 14.84x and 5.38x higher genotype missing rate than ForestQC, respectively (Figure S15a). In addition, ABHet has 2.09x (p-value = 2.183e-11) and VQSR has 2.13x higher genotype discordance rate (p-value = 1.584e-10) on than ForestQC (Figure S15b). For indels, ABHet and VQSR have 9.39x and 3.61x higher genotype missing rate on good indels than ForestQC, respectively (Figure S15c). Sample-level metrics also show that ForestQC has better Ti/Tv ratio on known SNVs (mean Ti/Tv: 1.75, 1.87, 1.82, 1.96 for No QC, ABHet, VQSR and ForestQC, respectively) and novel SNVs (mean Ti/Tv: 1.17, 1.03, 1.20, 1.39 for No QC, ABHet, VQSR and ForestQC, respectively) than other methods (Figures S15d and S15e). Paired t-tests for the difference in the mean Ti/Tv ratio of novel SNVs and known SNVs between ForestQC and other methods are all significant (p-value < 2.2e-16 versus all other methods). Similar to results of the BP dataset, ForestQC has higher accuracy in identifying good variants from gray variants, compared with ABHet approach and VQSR.

### Feature importance in random forest classifier

ForestQC uses several sequencing features in the random forest classifier to predict whether a variant with undermined quality is good or bad. To understand which sequencing features are more important indicators for quality of variants than other features, we analyze weight or importance of each feature that the random forest classifier learns during its model training. We first find that GC-content has the lowest importance in both BP and PSP datasets and also for both SNVs and indels (Figure S17). This means that GC-content may not be as a strong indicator of quality of variants as other features related to sequencing quality such as depth (DP) and genotype quality (GQ). Second, the results show that classification results are not determined by one or two most important features as there is no feature with much higher importance than other features except GC-content. This suggests that all sequencing features except GC-content are important indicators for quality of variants and need to be included in our model. We also check correlation among features and find that while certain pairs of features are highly correlated, like outlier GQ and mean GQ, SD DP and mean DP, some features have low correlation to other features, such as GC, suggesting that they may capture different information on quality of genetic variants (Figure S19). Third, we observe that the same features have different importance between the BP dataset and the PSP dataset. For example, for SNVs, an outlier ratio of GQ feature has the highest importance for the PSP dataset while it has the third lowest importance for the BP dataset (Figure S17a). Also, the importance of features varies between SNVs and indels. One example is a SD of DP feature that has the highest importance for SNVs in the BP dataset, but it has the third lowest importance for indels (Figure S17a, b). Therefore, these results suggest that each feature may have a different contribution to classification results depending on sequencing data and types of genetic variants.

### Performance of VQSR with different settings

For SNVs, GATK recommends three SNV call sets for training its VQSR model; 1) SNVs found in HapMap (“HapMap”), 2) SNVs in the omni genotyping array (“Omni”), and 3) SNVs in the 1000 Genomes Project (“1000G”). According to the VQSR parameter recommendation, SNVs in HapMap and Omni call sets are considered to contain only true variants while SNVs in 1000G contain both true and false positive variants(35). We call this recommended parameter setting “original VQSR.” We, however, find that considering SNVs in Omni to contain both true and false positive variants considerably improves the quality of good SNVs from VQSR for the BP dataset. We call this modified parameter setting “Omni_Modified VQSR”. Results show that the mean Ti/Tv on good novel SNVs from Omni_Modified VQSR is 1.76, which is much higher than that from original VQSR (1.41) and slightly higher than that from ForestQC (1.68) (Figure S19a). We also find that the mean number of total SNVs from Omni_Modified VQSR is 3.68M which is much smaller than that from original VQSR (3.99M) but higher than that from ForestQC (3.58M) (Figure S19b). In terms of other statistics, good SNVs from original VQSR has 3.66x higher ME rate, 7.40x higher genotype missing rate, and 1.16x higher genotype discordance rate (p-value = 0.0001118) than those SNVs from Omni_Modified VQSR (Figure S19c-e). Interestingly, we do not observe the improved performance of Omni_Modified VQSR for the PSP dataset as the mean novel Ti/Tv on good novel SNVs of Omni_Modified VQSR is 1.23, which is slightly smaller than that of original VQSR (1.27) (Figure S19a), although individuals have fewer good SNVs from Omni_Modified VQSR (3.53M) than that from original VQSR (3.75M) (Figure S19b). These results suggest that the performance of VQSR may change significantly depending on whether to consider a certain SNV call set to contain only true variants or both true and false positive variants, and it appears that the difference in performance is more noticeable in certain sequencing datasets than others.

Although Omni_Modified VQSR has slightly better Ti/Tv on good novel SNVs and identifies more good SNVs than does ForestQC, good SNVs from Omni_Modified VQSR have 2.76x higher ME rate, 13.20x higher genotype missing rate, and 1.35x higher genotype discordance rate (p-value < 2.2e-16) than good SNVs from ForestQC (Figure S19c-e). Hence, the results show that good SNVs from ForestQC have higher quality than those from VQSR even with the modification in the parameter setting.

## Discussion

We developed an accurate and efficient method called ForestQC to identify a set of variants with high sequencing quality from NGS data. ForestQC combines the traditional filtering approach for performing QC in GWAS and the classification approach that uses a machine learning algorithm to classify whether a variant has good quality. Our method first uses stringent filters to identify good and bad variants that unequivocally have high and low sequencing quality, respectively. ForestQC then trains a random forest classifier using the good and bad variants obtained from the filtering step, and predicts whether a variant with ambiguous quality (a gray variant) is good or bad in an unbiased manner. To evaluate ForestQC, we applied our method to two WGS datasets where one dataset consists of related individuals from families and the other dataset has unrelated individuals. We demonstrated that good variants identified from ForestQC in both datasets had higher sequencing quality than those from other approaches such as VQSR and a filtering approach based on ABHet.

To measure the performance of methods for variant quality control, one typically plans to apply these methods to benchmarking datasets where the true variants with high sequencing quality are verified. A few high-quality benchmarking variant sets have been proposed, including Genome In A Bottle (GIAB) (39), Platinum Genome (PlatGen) (40) and Syndip (41). GIAB has seven samples, PlatGen sequenced 17 individuals, and Syndip includes only two cell lines, CHM1 and CHM13. The sample sizes of these datasets are very small while we usually need to perform variant QC on an entire large dataset containing tens of millions of variants from hundreds of subjects or more. Thus, these datasets cannot be used as benchmarking datasets for variant QC. Apart, it is not expected to have a new benchmarking dataset with large sample size in the near future because it is expensive to construct such a dataset. Hence, in this study, we used real WGS datasets to evaluate different approaches for variant QC. Their large sample sizes allow more accurate calculation of various quality metrics and statistics used by the approaches for variant QC, and therefore enable more reliable performance evaluation.

To measure the quality of variants, we used 21 sample-level metrics and 20 variant-level metrics, plus genotype missing rate, ME rate and genotype discordance rate, resulting in a comprehensive evaluation of the performance of different methods. ME rate is found to be nearly linearly correlated with genotype errors(42–44), so it is a good quality metric for variants with pedigree information. Low genotype missing rate has been considered as an indicator of high-quality variant call set as a variant with high genotype missing rate indicates poor genotyping or sequencing quality(45). Also, high-quality variants would have the same genotypes generated by different genotyping technologies, such as sequencing and microarray. Thus, variant sequencing quality may be measured with genotype discordance rate between microarray and sequencing. One challenge with this approach is that genotypes generated by microarray are usually available for a small proportion of variants in the whole genome, especially for common and known variants, so it might not be able to show the sequencing quality of the entire variant call set. Another frequently used variant quality metric is Ti/Tv ratio (46–49). It is supposed to be around 2.0 for whole genome sequencing data(36). That is because transitions have higher frequency according to molecular mechanisms although the number of transversions is twice as many as transitions. Previous studies found that mitochondrial DNA and some non-human DNA sequences might be biased towards transitions or transversions(50, 51). In this study, we only computed Ti/Tv ratio for each QC method using the same human variant call set excluding mitochondria, in order to achieve an unbiased evaluation of all methods.

A main advantage of our approach over the traditional filtering approach is that our method does not attempt to classify gray variants using filters. It is difficult to determine the quality of those gray variants using filters if their QC metrics (e.g. genotype missing rate) are close to the thresholds of filters. Hence, ForestQC avoids a limitation of the traditional filtering approaches that determine the quality of every variant using filters, which may exclude some of good variants from the downstream analysis. We did not compare our approach with the traditional filtering approach used in GWAS that removes variants according to HWE p-values, ME rates and genotype missing rates. One main reason is that the performance of this approach changes dramatically depending on filters and thresholds for each filter, and there are numerous different thresholds of filters as well as many combinations of filters that could be tested. Another reason is that its performance could be arbitrarily determined depending on the filters we use. For example, if one filter is to remove any variants having more than zero Mendel errors, the ME rate of good variants would be zero, but we may be removing many other good variants. We checked the accuracy of a filtering approach based on ABHet as ABHet is often used in performing QC of NGS data and is a good indicator for variant quality(26,52,53). Also, as this approach is not based on standard QC metrics such as genotype missing rate, its performance is independent of those metrics unlike the standard filtering approaches. We showed that our approach outperformed the ABHet approach as the quality of good variants from ForestQC was better than that from ABHet, regardless of similar total number of good variants, as demonstrated by ME rate, missing rate, genotype discordance rate and Ti/Tv ratio in the BP and PSP dataset.

Although our approach is similar to VQSR as both approaches train machine learning classifiers to predict quality of variants, they have a few distinct differences. First, our approach trains the model using good and bad variants detected from sequencing data on which quality control is performed, while VQSR uses variants in existing databases, such as HapMap and 1000 genomes, as its training set. As VQSR uses previously known variants for model training, good variants from VQSR are likely to contain more known (and likely to be common) variants than novel (and rare) variants. We showed in both WGS datasets that it did identify more common and known SNVs and indels as good variants than ForestQC. This may not be a desirable outcome for some sequencing studies if one of their main goals is to identify rare and novel variants not captured in chips. Another difference between ForestQC and VQSR is the set of features used in the classifiers. While both methods use features related to sequencing depth and genotyping quality, VQSR uses some features that are specifically calculated by GATK software while our method uses quality information reported in the standard VCF file. This suggests that our method is more generalizable than VQSR as it can be applied to VCF files generated from variant callers other than GATK. The last difference is the machine learning algorithms that ForestQC and VQSR use. Our method trains a random forest model while VQSR trains a Gaussian Mixture model. Using the BP and PSP dataset, we found that random forest model was much faster than Gaussian Mixture model (Table S11).

In addition to SNVs, we applied our method to indels in both WGS datasets and found that indels had much lower sequencing quality than do SNVs as the fraction of good indels detected by ForestQC was considerably smaller than that of SNVs. This is somewhat expected because indel or structural variant calling is much more difficult than SNV calling from sequencing data, and some of them are likely to be false positives(54, 55). It is, however, important to note that VQSR classifies many more indels as good variants than does ForestQC or ABHet, but those good indels from VQSR may not have high sequencing quality. We showed that good indels from VQSR had similar Mendelian error rate to that without performing QC, indicating the poor performance of VQSR on indels. VQSR considers indels from Mills gold standard call set(37) as true variants, and while those indels might represent true variant sites, it does not necessarily mean that genotyping on those sites is accurate. Therefore, genetic studies need to perform stringent QC on indels to remove those erroneous calls and not to have false positive findings in their downstream analysis.

We found that the performance of VQSR was improved dramatically for the BP dataset when we considered SNVs in Omni genotyping array to have both true and false positive sites, compared with when they were assumed to have all true sites. We, however, did not observe this performance enhancement for the PSP dataset. This suggests that users may need to try different parameter settings to obtain optimal results from VQSR for specific sequencing datasets they analyze. Another issue with VQSR and also with ABHet is that some of good SNVs or indels have high genotype missing rate and ME rate, which may not be suitable for the downstream analysis such as association analysis. Thus, those variants need to be filtered out separately, which means users may need to perform an additional filtering step in addition to applying VQSR and ABHet to the dataset. As the filtering step is incorporated in ForestQC, our method does not have this issue.

Our approach is an extension of a previous approach that uses a logistic regression model to predict the quality of variants in the BP dataset(30). While our approach is similar to the previous approach in that they both combine filtering and classification approaches, ForestQC uses a random forest classifier that has higher accuracy than a logistic regression model, according to our simulation results. It includes more bad variants for model training, leading to predictions with fewer biases. ForestQC also includes more features than the previous approach as well as more filters to improve the quality of good variants. Additionally, compared with the previous approach, ForestQC is more user-friendly and generalizable because users can choose or define different features and filters and tune the parameters according to their research goals.

ForestQC is efficient, modularized and flexible with following features. First, users are allowed to change thresholds for filters as needed. This is important because filters that are stringent for one dataset may not be stringent for another dataset. For example, variants from sequence data with very small sample size (e.g. < 100) may not have statistical power to have significant HWE p-values, and hence higher p-value thresholds may need to be used, compared with studies with larger sample size. If filters are not stringent enough, there may be many bad variants, and ForestQC would train a very stringent classifier, leading to the possible removal of good variants. On the contrary, if the filters are too stringent, there would be too few good variants or bad variants, which would lower the accuracy of our random forest classifier. In this study, after the filtering step, 4.39% of SNVs and 15.72% of indels in the BP dataset, and 5.06% of SNVs and 15.66% of indels in PSP dataset, were determined as bad variants. Empirically, we suggest filters for ForestQC such that after the filtering step, a fraction of bad variants is about 4-16%. Normally, the default parameter settings are recommended, which are the same sets of filters and features described in this paper. The selection of threshold values for these filters are based on our previous study for WGS data of extended pedigrees for bipolar disorder(30). Second, users are allowed to use their own filters and features provided that they specify values for those new filters and features at each variant site, and our software also allows users to remove existing filters and features. As there may be filters and features that capture sequencing quality of variants more accurately than current set of filters and features, this option allows users to improve ForestQC further. For example, users can employ mappability, strand bias and micro-repeats as features, instead of sequencing depth and genotyping quality used in this study, because DP and GQ might penalize disease-causing variants with low coverage. Also, if users want to obtain more variants after QC, they may lower the standard for good variants, that is, increase the threshold values of ME or missing rate for determining good variants. Third, ForestQC generates the probability of each gray variant being a good variant. This probability needs to be greater than a certain threshold for a gray variant to be predicted to be good, and it can also be used to analyze sequencing quality of certain variants. If studies find that a certain gray variant is associated with a phenotype, they may consider checking whether its probability of being a good variant is high enough. Lastly, ForestQC allows users to change the probability threshold for determining whether each gray variant is good or bad. Users may lower this threshold if they are interested in obtaining more good variants at the cost of including more bad variants.

## Materials and Methods

### ForestQC

ForestQC consists of two approaches: a filtering approach and a machine learning approach based on a random forest algorithm.

#### Filtering

Given a variant call set from next generation sequencing data, ForestQC first applies several stringent filters to identify good, bad, and gray variants. Good variants are ones that pass all filters while bad variants fail any of them (Table S2, S3). Gray variants are variants that neither pass filters for good variants nor fail filters for bad variants. We use following filters in the filtering step.

- Mendelian error (ME) rate. The Mendelian error occurs when a child’s genotype is inconsistent with genotypes from parents. ME rate is calculated as the number of ME among all trios divided by the number of trios for a given variant. Note that this statistic is only available for family-based data.
- Genotype missing rate. This is the proportion of missing alleles in each variant.
- Hardy-Weinberg equilibrium (HWE) p-value. This is a p-value for hypothesis testing whether a variant is in Hardy-Weinberg equilibrium. Its null hypothesis is that the variant is in Hardy-Weinberg equilibrium. We use the algorithm used in an open-source software, VCFtools(56) for the calculation of Hardy-Weinberg equilibrium p-value.
- ABHet. This is allele balance for heterozygous calls. ABHet is calculated as the number of reference reads from individuals with heterozygous genotypes divided by the total number of reads from such individuals, which is supposed to be 0.50 for good variants. For variants in chromosome X, we only calculate ABHet for females.

#### Random forest classifier

Random forest algorithm is a machine learning algorithm that runs efficiently on large datasets with high accuracy(34). Briefly, random forest builds several randomized decision trees, each of which is trained to classify the input objects. For classification of a new object, the fitted random forest model passes the input vector down to each of the decision trees in the forest. Each decision tree has its classification result, then the forest would output the classification that the majority of the decision trees make. Balancing efficiency and accuracy, we train a random forest classifier using 50 decision trees (Figure S2) and 50% as probability threshold (Figure S3).

To train random forest, we use good and bad variants identified from the previous filtering step as a training dataset, after balancing their sample size by random sampling. Normally, good variants are much more numerous than bad variants, so we randomly sample from good variants with the sample size of bad variants. Hence, the sample size of the balanced training set would be twice as large as the sample size of bad variants. We also need features in training random forest, which characterize datasets, and we use following features.

- Mean and standard deviation of depth (DP) and genotyping quality (GQ). Depth and genotyping quality values are extracted from DP and GQ fields of each sample in VCF files, respectively, and mean and standard deviation are calculated over all samples for each variant.
- Outlier depth and outlier genotype quality. These are the proportions of samples whose DP or GQ is lower than a particular threshold. We choose this threshold as the first quartile value of all DP or GQ values of variants on chromosome 1. We use DP and GQ of variants on only chromosome 1 to reduce the computational costs.
- GC content: We first split a reference genome into window size of 1,000 bp and calculate GC content for each window as (# of G or C alleles) / (# of A, G, C or T alleles). Then, each variant is assigned a GC content value according to its position in the reference genome.

After training random forest with the training dataset using above features, we next use the fitted model to make predictions on gray variants on being good variants. Gray variants with the predicted probability of being good larger than 50% are labeled as predicted good variants. Then the predicted good variants and good variants from the previous filtering step are combined to form the final set of good variants. We apply the same procedure to identify bad variants.

### Comparison of different machine learning algorithms

We compare eight different machine learning algorithms, in order to identify the best algorithm used for ForestQC. They are 1) k-nearest neighbors for supervised 2-class classification (8 threads); 2) logistic regression (8 threads); 3) single support vector machine with Gaussian kernel function and penalty parameter C of 1 (1 thread); 4) random forest with 50 trees (8 threads); 5) naïve Bayes without any prior probabilities of the classes (1 thread); 6) artificial neural network with sigmoid function as activation function (8 threads). It has 1 hidden layer with 10 units; 7) AdaBoost with 50 estimators and learning rate of 1, which uses SAMME.R real boosting algorithm (1 thread); 8) and quadratic discriminant analysis without any prior on classes. Its regularization is 0 and its threshold for rank estimation is 1e-4 (1 thread). Other parameters of these machine learning algorithm are default, as described in the documentations of Python scikit-learn package(57). All learning algorithms use the seven aforementioned features: mean and standard deviation of sequencing depth, mean and standard deviation of genotype quality, outlier depth, outlier quality and GC content.

To test these eight machine learning algorithms, we obtain training and test datasets from the BP dataset, using filters described in Table S2 and S3. There are 21,248,103 good SNVs and 2,257,506 good indels while there are 1,100,325 bad SNVs and 624,965 bad indels. We sample 100,000 variants randomly from good variants and 100,000 variants from bad variants to generate a training set. Similarly, 100,000 good variants and 100,000 bad variants are randomly chosen from the rest of variants to form a test set. Each machine learning model shares the same training and test sets. We train the machine learning models and measure training time at a training stage, and then test their accuracy and measure prediction time at a testing stage. We measure the time cost of each algorithm, which is the elapsed clock time between the start and end of each algorithm. To assess the performance of each algorithm, we compute F1-score for the test set. F1-score is the harmonic average of precision and recall, which is calculated as 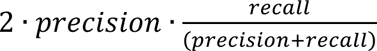. The closer F1-score is to 1, the higher classification accuracy is. Recall is the fraction of true positive results over all samples that should be given positive prediction. Precision is the number of true positive results divided by the number of positive results predicted by the classifier. We also measure the model accuracy using 10-fold cross validation, as well as the area under the receiver operating characteristic curve.

### ABHet approach and VQSR

We compare ForestQC with two other approaches for performing QC on genetic variants. One is a filtering approach based on ABHet and the other is a classification approach called VQSR from GATK software. For the ABHet approach, we consider variants with ABHet > 0.7 or < 0.3 as bad variants, and the rest as good variants. We chose this threshold setting of ABHet (> 0.3 and < 0.7) because the ADSP project could not reliably confirm heterozygous calls with ABHet > 0.7 with Sanger sequencing(26). We also exclude variants with small ABHet values (< 0.3) to ensure high quality. For GATK, we use recommended arguments as of 2018-04-04(35). For SNVs, VQSR takes SNVs in HapMap 3 release 3, 1000 Genome Project and Omni genotyping array as training resources, and dbSNP135 as known sites resource. HapMap and Omni sites are considered as true sites, meaning that SNVs in these datasets are all true variants, while 1000 Genome Project sites are regarded as false sites, meaning that there could be both true and false-positive variants. The desired level of sensitivity of true sites is set to be 99.5%. In the BP dataset, we run VQSR version 3.5-0-g36282e4 with following annotations; quality by depth (QD), RMS mapping quality (MQ), mapping quality rank sum test (MQRankSum), read position rank sum test (ReadPosRankSum), fisher strand (FS), coverage (DP) and strand odds ratio (SOR) to evaluate the likelihood of true positive calls. In the PSP dataset, we use VQSR version 3.2-2-gec30cee that uses all previous annotations and additional inbreeding coefficient (InbreedingCoeff) except SOR because variants in PSP dataset do not have the SOR annotation. For indels, VQSR takes indels in Mills gold standard call set(37) as true training resource, and dbSNP135 as known sites resource. The desired level of sensitivity of true sites is set to be 99.0%. We use VQSR version 3.5-0-g36282e4 with QD, DP, FS, SOR, ReadPosRankSum and MQRankSum annotations to evaluate the likelihood of true positive calls in the BP dataset, while we run VQSR version 3.2-2-gec30cee with the same annotations and additional InbreedingCoeff except SOR for the PSP dataset.

### BP and PSP WGS datasets

The BP WGS dataset is for studying bipolar disorder whose average coverage is 36. This study recruited individuals from 11 Colombia (CO) and 15 Costa Rica (CR) extended pedigrees in total. 454 subjects from 10 CO and 12 CR families are both whole genome sequenced and genotyped with microarray. There are 144 individuals diagnosed with BP1 and 310 control samples that are unaffected or have non-BP traits. We use highly scalable Churchill pipeline(58) to do variant calling for the BP data set, where GATK-HaplotypeCaller 3.5-0-g36282e4 is used as the variant caller according to the GATK best practices(23) and the reference genome is HG19. After initial QC on individuals, five individuals are removed because of poor sequencing quality and possible sample mix-ups. Finally, 449 individuals are included in an analysis, resulting in 25,081,636 SNVs and 3,976,710 indels. 1,814,326 SNVs in the WGS dataset are also genotyped with microarray, which are used to calculate genotype discordance rate. In this study, we use the BP dataset before any QC performed on genetic variants. In a previous study(30), genetic variants in the BP WGS dataset are first processed with VQSR and then filtered with a trained logistic regression model to remove variants with low quality.

The PSP WGS dataset is for studying progressive supranuclear palsy with average coverage of 29. 544 unrelated individuals are whole genome sequenced, 518 of whom are also genotyped with microarray. Among them, 119 individuals have 547,644 SNPs and 399 individuals have 1,682,489 SNPs genotyped with microarray, respectively. That 119 individuals would be excluded when calculating genotype discordance rate in case of biases caused by fewer SNPs. There are 356 individuals diagnosed with PSP and 188 individuals as controls. Variant calling for the PSP dataset is performed using Churchill pipeline, where GATK-HaplotypeCaller 3.2-2-gec30cee is used as the variant caller according to the GATK best practices and the reference genome is HG19. 49 samples are found to have high missing rate or high relatedness with other samples, or are diagnosed with diseases other than PSP, so they are removed. Next, we extract variant data with only 495 individuals with VCFtools. Monomorphic variants are then removed. After preprocessing, the PSP WGS dataset has 33,273,111 SNVs and 5,093,443 indels. There are 1,682,489 SNVs from 381 samples genotyped by both microarray and WGS, which are used for calculating genotype discordance rate.

### Performance metrics

21 sample-level metrics and 20 variant-level metrics are defined to measure the sequencing quality of the variant call set after performing quality control (Table S12). Note that we do not show all sample-level metrics and variant-level metrics in the main text. Other metrics are available in supplemental materials. Variant-level metrics provide us with a summarized assessment report of the sequencing quality of a variant call set, such as total SNVs of the whole dataset. They are calculated based on the information of all variants in a variant call set. For example, the number and the proportion of multi-allelic SNVs are counted for the entire dataset, each of which is identified according to its reference and alternate alleles. On the other hand, sample-level metrics enable the inspection of the sequencing quality for sequenced individuals in a variant call set. For instance, we check the distribution of novel Ti/Tv or other quality metrics among all individuals in the study. Sample-level metrics are calculated for each sample, using its genotype information on all variants in the dataset, and a distribution of those metrics across all individuals is shown as a box plot. For example, the number of SNV singletons on a sample level shows the distribution of the number of SNV singletons across all sequenced individuals. In this study, both sample-level and variant-level metrics are used to evaluate the sequencing quality of WGS variant datasets.

In addition, we also use genotype missing rate, ME rate and genotype discordance rate as variant quality metrics, which are computed using the entire variant call set. The definitions of genotype missing rate and ME rate have been described above. Note that ME rate is only available for family-based datasets, such as the BP dataset, so we do not calculate ME rate for the PSP dataset that only includes unrelated individuals. Genotype discordance rate is the proportion of individuals whose genotypes are inconsistent between next-generation sequencing and microarray. This metric can only be calculated with a subset of variants due to the limited number of variants genotyped by both sequencing and microarray. Note that microarray might also have biases in genotyping, leading to some limitations of genotype discordance rate. For example, microarray usually genotype selected variants, especially common and known variants, so genotype discordance rate is only available for these selected variants and it cannot provide quality evaluation for all variants, especially rare variants. Genotype missing rate, ME rate and genotype discordance rate provide us with accurate evaluation of variant quality, because true positive variants with high quality are very likely to have low values of these three metrics.

## Acknowledgements

We thank Dr. Susan K. Service from Department of Psychiatry and Biobehavioral Sciences, UCLA for the precious comments and suggestions to our project and this manuscript. We thank all study participants in the BP and PSP datasets.

## Availability and Implementation

ForestQC is available at https://github.com/avallonking/ForestQC under the open source MIT license. There are detailed installation instructions and user guide. It is implemented with Python3 and is compatible with Linux, Mac OSX and Windows 64-bit operating systems.

## Authors’ contributions

JL and JHS designed the method and conceived this study. JL developed the method and did the analysis of the BP and PSP datasets. JHS did preprocessing and variant calling for the BP dataset. BJ and SH did preprocessing and variant calling for the PSP dataset. LZ tested the software. NF provided the BP dataset and GC contributed the PSP dataset. JL and JHS wrote the manuscript. All authors read and approved the final manuscript.

## Supporting Information

Supporting Information includes 19 figures and 11 tables. Captions listed below.

Figure S1: Receiver operating charateristic (ROC) curves and area under the curve of eight machine learning models.

Figure S2: Relationship between the number of trees in random forest model and the performance of ForestQC. Relationship between the number of trees and (a) CPU time and (b) F1-score.

Figure S3: Relationship between the probability threshold for predicting a variant to be good and the precision of ForestQC. If the probability of a variant predicted to be good is larger than the probability threshold, this variant would be labeled as a good variant. Classification precision changes along with the probability threshold in SNV classification (a) and indel classification (b). The precision of ForestQC is measured in F1-score.

Figure S4: Overall quality of good and bad variants in the BP dataset identified by ForestQC using ME rate as a filter or not. The average Mendelian error rate and genotype missing rate for SNVs and indels, and genotype discordance rate to microarray data for SNVs are shown. Data are represented as the mean ± SEM.

Figure S5: Sample-level quality metrics of good variants in the BP dataset identified by ForestQC using ME rate as a filter or not. (a) Total number of SNVs. (b) The number of SNVs found in dbSNP. (c) the number of SNVs not found in dbSNP. (d) Ti/Tv ratio of SNVs found in dbSNP. (e) Ti/Tv ratio of SNVs not found in dbSNP. (f) Total number of indels. (g) the number of indels found in dbSNP. (h) the number of indels not found in dbSNP. The version of dbSNP is 150.

Figure S6: Overall quality of bad variants in the BP dataset detected by four different methods, including no QC applied, ABHet approach, VQSR and ForestQC. The average Mendelian error rate and genotype missing rate for SNVs and indels, and genotype discordance rate to microarray data for SNVs are shown. Data are represented as the mean ± SEM.

Figure S7: Sample-level quality metrics of good variants in the BP dataset identified by four different methods, including no QC applied, ABHet approach, VQSR and ForestQC. (a) Ti/Tv ratio of SNVs found in dbSNP. (b) The number of SNVs found in dbSNP. (c) The number of indels found in dbSNP. (d) The number of indels not found in dbSNP. The version of dbSNP is 150.

Figure S8: Overall quality of rare variants (MAF < 0.03) and common variants (MAF 0.03) in the BP dataset. The average Mendelian error rate and genotype missing rate for SNVs and indels, and genotype discordance rate to microarray data for SNVs are shown. Data are represented as the mean ± SEM.

Figure S9: Overall quality of good variants identified from gray variants in the BP dataset processed by four different methods, including no QC applied, ABHet approach, VQSR and ForestQC. The average Mendelian error rate and genotype missing rate for SNVs and indels, and genotype discordance rate to microarray data for SNVs are shown. Data are represented as the mean ± SEM.

Figure S10: Sample-level quality metrics of good variants identified from gray variants in the BP dataset processed by four different methods, including no QC applied, ABHet approach, VQSR and ForestQC. (a) Total number of SNVs. (b) The number of SNVs found in dbSNP. (c) the number of SNVs not found in dbSNP. (d) Ti/Tv ratio of SNVs found in dbSNP. (e) Ti/Tv ratio of SNVs not found in dbSNP. (f) Total number of indels. (g) the number of indels found in dbSNP. (h) the number of indels not found in dbSNP. The version of dbSNP is 150.

Figure S11: Selected sample-level quality metrics of good variants in BP dataset identified by VQSR using “SOR” or not. (a) Ti/Tv ratio of SNVs not found in dbSNP, (b) the number of total SNVs and (c) the number of total indels in the BP dataset processed with VQSR using “SOR” or not. SOR stands for StrandOddsRatio, which is a metric for strand bias measured by the Symmetric Odds Ratio test. The version of dbSNP is 150.

Figure S12: Overall quality of bad variants in the PSP dataset detected by four different methods, including no QC applied, ABHet approach, VQSR and ForestQC. The average genotype missing rate for both SNVs and indels, and genotype discordance rate to microarray data for SNVs are shown. Data are represented as the mean ± SEM.

Figure S13: Sample-level quality metrics of good variants in PSP dataset identified by four different methods, including no QC applied, ABHet approach, VQSR and ForestQC. (a) Ti/Tv ratio of SNVs found in dbSNP. (b) The number of SNVs found in dbSNP. (c) The number of indels found in dbSNP. (d) The number of indels not found in dbSNP. The version of dbSNP is 150.

Figure S14: Overall quality of rare variants (MAF < 0.03) and common variants (MAF 0.03) in the PSP dataset. The average genotype missing rate for SNVs and indels, and genotype discordance rate to microarray data for SNVs are shown. Data are represented as the mean ± SEM.

Figure S15: Overall quality of good variants identified from gray variants in the PSP dataset processed by four different methods, including no QC applied, ABHet approach, VQSR and ForestQC. The average genotype missing rate for both SNVs and indels, and genotype discordance rate to microarray data for SNVs are shown. Data are represented as the mean ± SEM.

Figure S16: Sample-level quality metrics of good variants identified from gray variants in the PSP dataset processed by four different methods, including no QC applied, ABHet approach, VQSR and ForestQC. (a) Total number of SNVs. (b) The number of SNVs found in dbSNP. (c) the number of SNVs not found in dbSNP. (d) Ti/Tv ratio of SNVs found in dbSNP. (e) Ti/Tv ratio of SNVs not found in dbSNP. (f) Total number of indels. (g) the number of indels found in dbSNP. (h) the number of indels not found in dbSNP. The version of dbSNP is 150.

Figure S17: Feature importance of each feature in the random forest model of ForestQC applied to the BP and PSP datasets. DP stands for sequencing depth. GQ stands for genotyping quality. SD means standard deviation. Outlier DP or GQ means the proportion of samples having genotyping quality or sequencing depth lower than the first quartile of depth or genotyping quality in chromosome 1. GC stands for the GC content of a 1000-bp window where the variant is located. (a) Feature importance in SNV classification. (b) Feature importance in indel classification.

Figure S18: Pearson’s correlation coefficients between each pair of features in the BP and PSP dataset.

Figure S19: Quality of good SNVs identified by VQSR with two different settings of training resources and ForestQC. (a) Ti/Tv ratio of SNVs not found in dbSNP v150 and (b) total number of SNVs in the BP and PSP dataset. (c)-(e) Average Mendelian error rate, average genotype missing rate, and average genotype discordance rate of good SNVs in the BP dataset. Data are represented as the mean ± SEM. “Omni_Modified VQSR”: SNVs in Omni chip array call set are considered to contain both true and false positive sites. “original VQSR”: SNVs in Omni chip array call set are considered to contain only true sites.

Table S1: Accuracy of eight different machine learning algorithms

Table S2: Thresholds of four filters for the selection of good variants from the original dataset

Table S3: Thresholds of four filters for the selection of bad variants from the original dataset

Table S4: Variant-level quality metrics of variants in the BP dataset processed by ForestQC with different settings

Table S5: Variant-level quality metrics of good variants in the BP dataset processed by different methods

Table S6: Rare variants and common variants in the BP dataset processed by different methods

Table S7: Variant-level quality metrics of good variants identified from gray variants in the BP dataset

Table S8: Variant-level quality metrics of good variants in the PSP dataset processed by four different methods

Table S9: Rare variants and common variants in the PSP dataset processed by different methods

Table S10: Variant-level quality metrics of good variants identified from gray variants in the PSP dataset

Table S11: Running time of ForestQC and VQSR in two datasets, measured in real time

Table S12: Definitions of 23 metrics for sequencing quality control calculated for sample-level and variant-level

