## Supporting information for "ForestQC: quality control on genetic variants from next-generation sequencing data using random forest"

Fig S1: Receiver operating charateristic (ROC) curves and area under the curve (AUC) of eight machine learning models in (a) SNV classification and (b) indel classification.

Fig S2: Relationship between the number of trees in random forest model and the performance of ForestQC. Relationship between the number of trees and (a) CPU time and (b) F1-score.

Fig S3: Relationship between the probability threshold for predicting a variant to be good and the precision of ForestQC. If the probability of a variant predicted to be good is larger than the probability threshold, this variant would be labeled as a good variant. Classification precision changes along with the probability threshold in SNV classification (a) and indel classification (b). The precision of ForestQC is measured in F1-score.

Fig S4: Overall quality of good and bad variants in the BP dataset identified by ForestQC using ME rate as a filter or not. The average Mendelian error rate and genotype missing rate for SNVs and indels, and genotype discordance rate to microarray data for SNVs are shown. Data are represented as the mean ± SEM.

Fig S5: Sample-level quality metrics of good variants in the BP dataset identified by ForestQC using ME rate as a filter or not. (a) Total number of SNVs. (b) The number of SNVs found in dbSNP. (c) the number of SNVs not found in dbSNP. (d) Ti/Tv ratio of SNVs found in dbSNP. (e) Ti/Tv ratio of SNVs not found in dbSNP. (f) Total number of indels. (g) the number of indels found in dbSNP. (h) the number of indels not found in dbSNP. The version of dbSNP is 150.

Fig S6: Overall quality of bad variants in the BP dataset detected by four different methods, including no QC applied, ABHet approach, VQSR and ForestQC. The average Mendelian error rate and genotype missing rate for SNVs and indels, and genotype discordance rate to microarray data for SNVs are shown. Data are represented as the mean ± SEM.

Fig S7: Sample-level quality metrics of good variants in the BP dataset identified by four different methods, including no QC applied, ABHet approach, VQSR and ForestQC. (a) Ti/Tv ratio of SNVs found in dbSNP. (b) The number of SNVs found in dbSNP. (c) The number of indels found in dbSNP. (d) The number of indels not found in dbSNP. The version of dbSNP is 150.

Fig S8: Overall quality of rare variants (MAF < 0.03) and common variants (MAF $\geq$ 0.03) in the BP dataset. The average Mendelian error rate and genotype missing rate for SNVs and indels, and genotype discordance rate to microarray data for SNVs are shown. Data are represented as the mean ± SEM.

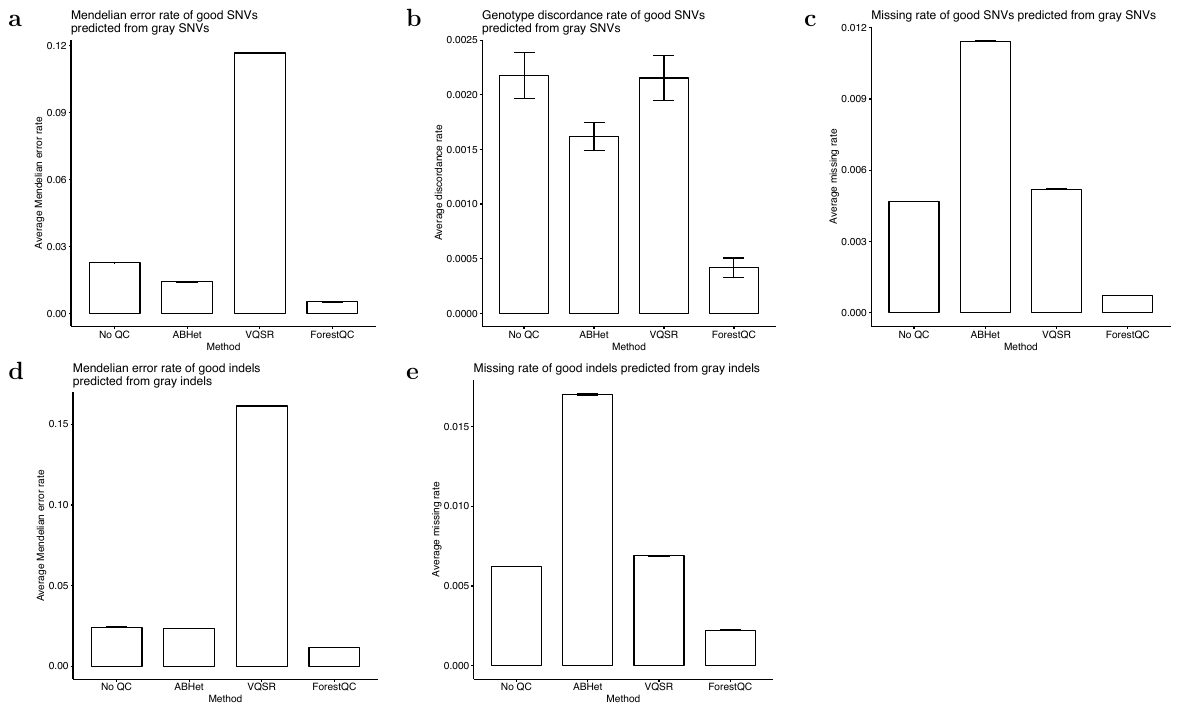

Fig S9: Overall quality of good variants identified from gray variants in the BP dataset processed by four different methods, including no QC applied, ABHet approach, VQSR and ForestQC. The average Mendelian error rate and genotype missing rate for SNVs and indels, and genotype discordance rate to microarray data for SNVs are shown. Data are represented as the mean ± SEM.

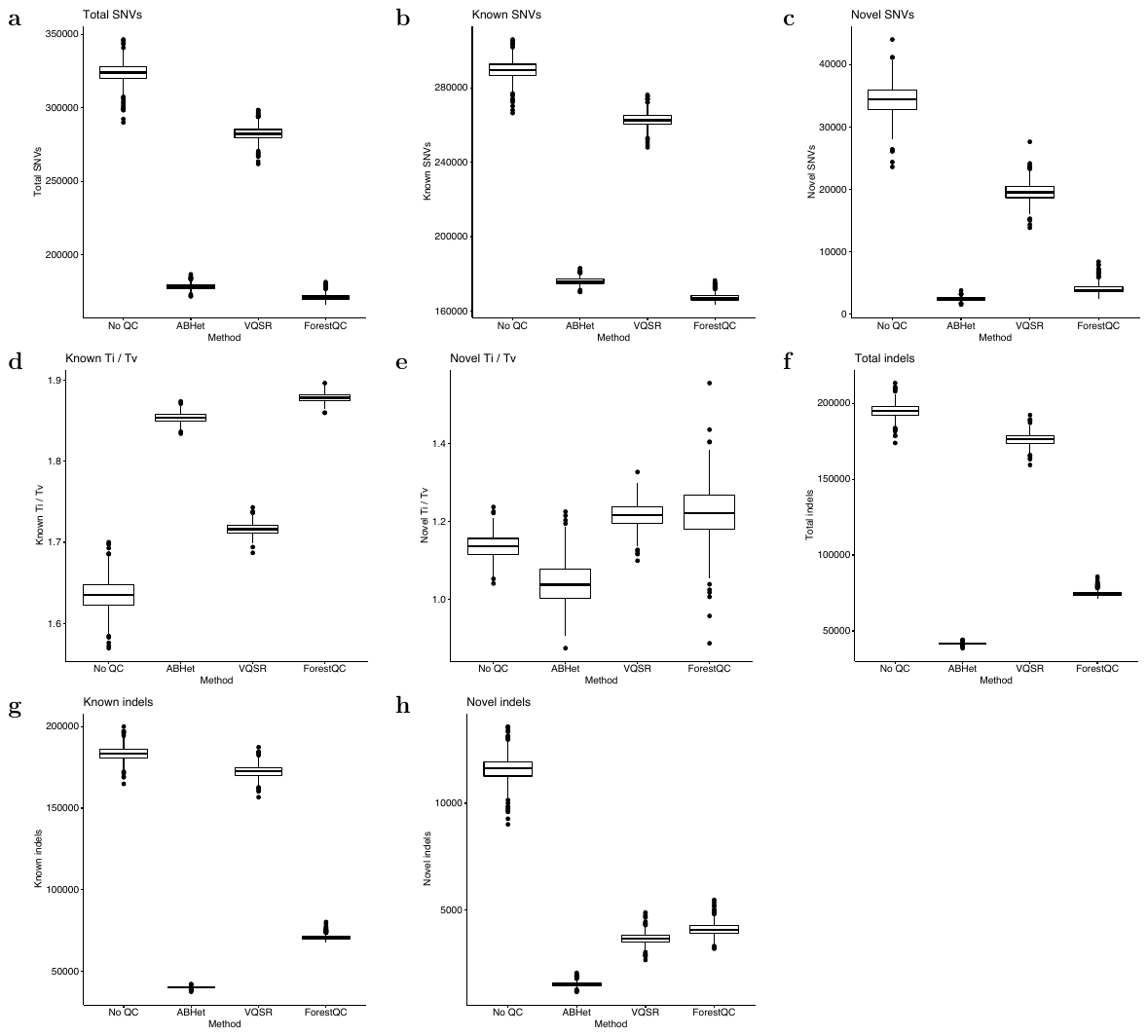

Fig S10: Sample-level quality metrics of good variants identified from gray variants in the BP dataset processed by four different methods, including no QC applied, ABHet approach, VQSR and ForestQC. (a) Total number of SNVs. (b) The number of SNVs found in dbSNP. (c) the number of SNVs not found in dbSNP. (d) Ti/Tv ratio of SNVs found in dbSNP. (e) Ti/Tv ratio of SNVs not found in dbSNP. (f) Total number of indels. (g) the number of indels found in dbSNP. (h) the number of indels not found in dbSNP. The version of dbSNP is 150.

Fig S11: Selected sample-level quality metrics of good variants in BP dataset identified by VQSR using “SOR” or not. (a) Ti/Tv ratio of SNVs not found in dbSNP, (b) the number of total SNVs and (c) the number of total indels in the BP dataset processed with VQSR using “SOR” or not. SOR stands for StrandOddsRatio, which is a metric for strand bias measured by the Symmetric Odds Ratio test. The version of dbSNP is 150.

Fig S12: Overall quality of bad variants in the PSP dataset detected by four different methods, including no QC applied, ABHet approach, VQSR and ForestQC. The average genotype missing rate for both SNVs and indels, and genotype discordance rate to microarray data for SNVs are shown. Data are represented as the mean ± SEM.

Fig S13: Sample-level quality metrics of good variants in PSP dataset identified by four different methods, including no QC applied, ABHet approach, VQSR and ForestQC. (a) Ti/Tv ratio of SNVs found in dbSNP. (b) The number of SNVs found in dbSNP. (c) The number of indels found in dbSNP. (d) The number of indels not found in dbSNP. The version of dbSNP is 150.

Fig S14: Overall quality of rare variants (MAF < 0.03) and common variants (MAF $\geq$ 0.03) in the PSP dataset. The average genotype missing rate for SNVs and indels, and genotype discordance rate to microarray data for SNVs are shown. Data are represented as the mean ± SEM.

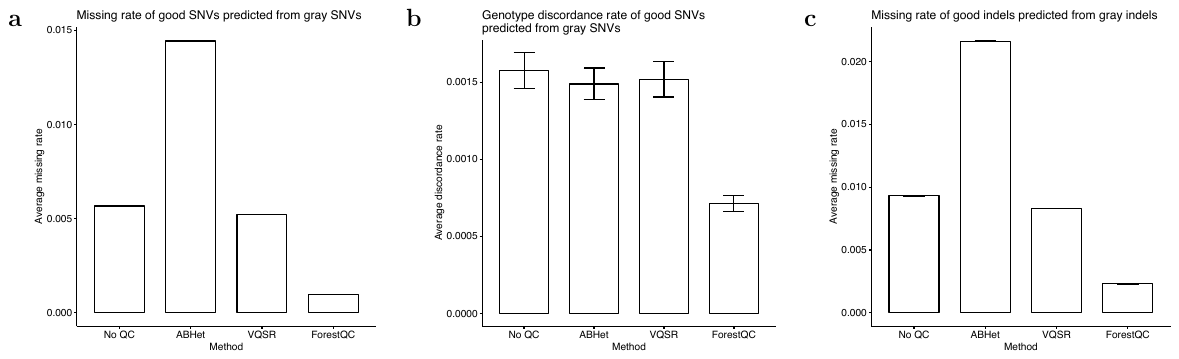

Fig S15: Overall quality of good variants identified from gray variants in the PSP dataset processed by four different methods, including no QC applied, ABHet approach, VQSR and ForestQC. The average genotype missing rate for both SNVs and indels, and genotype discordance rate to microarray data for SNVs are shown. Data are represented as the mean ± SEM.

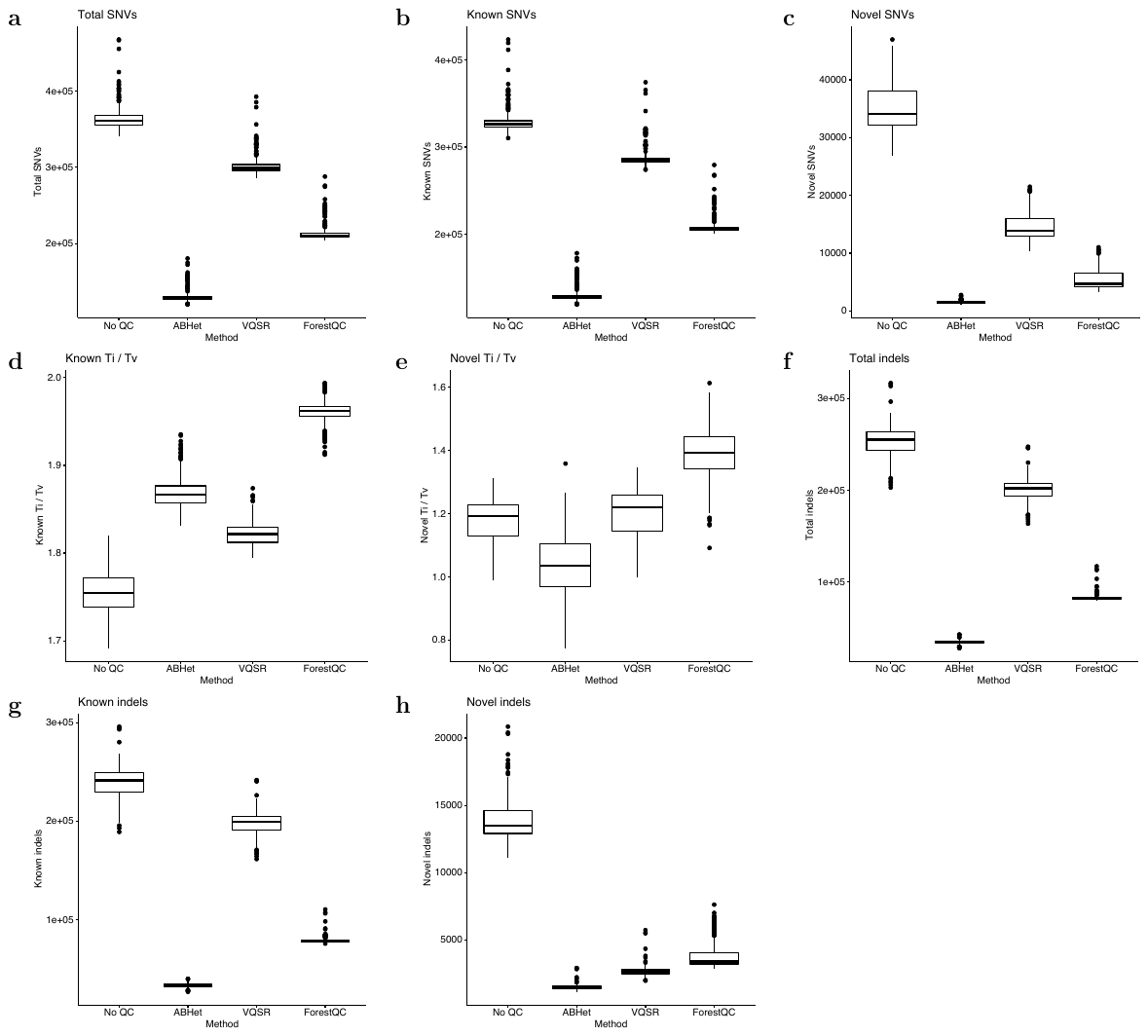

Fig S16: Sample-level quality metrics of good variants identified from gray variants in the PSP dataset processed by four different methods, including no QC applied, ABHet approach, VQSR and ForestQC. (a) Total number of SNVs. (b) The number of SNVs found in dbSNP. (c) the number of SNVs not found in dbSNP. (d) Ti/Tv ratio of SNVs found in dbSNP. (e) Ti/Tv ratio of SNVs not found in dbSNP. (f) Total number of indels. (g) the number of indels found in dbSNP. (h) the number of indels not found in dbSNP. The version of dbSNP is 150.

Fig S17: Feature importance of each feature in the random forest model of ForestQC applied to the BP and PSP datasets. DP stands for sequencing depth. GQ stands for genotyping quality. SD means standard deviation. Outlier DP or GQ means the proportion of samples having genotyping quality or sequencing depth lower than the first quartile of depth or genotyping quality in chromosome 1. GC stands for the GC content of a 1000-bp window where the variant is located. (a) Feature importance in SNV classification. (b) Feature importance in indel classification.

Fig S18: Pearson’s correlation coefficients between each pair of features in the BP and PSP dataset.

Fig S19: Quality of good SNVs identified by VQSR with two different settings of training resources and ForestQC. (a) Ti/Tv ratio of SNVs not found in dbSNP v150 and (b) total number of SNVs in the BP and PSP dataset. (c)-(e) Average Mendelian error rate, average genotype missing rate, and average genotype discordance rate of good SNVs in the BP dataset. Data are represented as the mean ± SEM. “Omni_Modified VQSR”: SNVs in Omni chip array call set are considered to contain both true and false positive sites. “original VQSR”: SNVs in Omni chip array call set are considered to contain only true sites.

**Table S1: Thresholds of four filters for the selection of good variants from the original dataset**

| Filter | Threshold |
| --- | --- |
| Mendelian error rate | 0 |
| Missing rate | < 0.5% |
| HWE p-value | > 0.01 |
| ABHet | 0.3 – 0.7 |

Each good variant must satisfy all thresholds. “HWE p-value”: p-value in testing for Hardy-Weinberg equilibrium.

**Table S2: Thresholds of four filters for the selection of bad variants from the original dataset**

| Condition | Filter | Rare variants  (MAF < 0.03) | Common variants  (MAF $\geq$ 0.03) |
| --- | --- | --- | --- |
| ALL | Mendelian error rate | > 3 / (# of trios) | > 5 / (# of trios) |
|  | Missing rate | > 2% | > 3% |
|  | HWE p-value | < 0.005 | < 0.0005 |
|  | ABHet | > 0.75 or < 0.25 | > 0.75 or < 0.25 |
| ANY | Mendelian error rate | > 8 / (# of trios) | > 10 / (# of trios) |
|  | Missing rate | > 8% | > 10% |
|  | HWE p-value | < 0.001 | < 1e-8 |

ALL means all thresholds should be satisfied. ANY means variants are considered bad if they satisfy any one of the thresholds. Note that rare variants (MAF < 0.03) and common variants (MAF $\geq$ 0.03) have different thresholds. “HWE p-value”: p-value in testing for Hardy-Weinberg equilibrium.

**Table S3: Accuracy of eight different machine learning algorithms**

| Machine learning algorithm | Accuracy in  SNV classification | Accuracy in  indel classfication |
| --- | --- | --- |
| Random Forest | 0.9736 $\pm$ 0.0018 | 0.9428 $\pm$ 0.0024 |
| ANN | 0.9707 $\pm$ 0.0016 | 0.9401 $\pm$ 0.0027 |
| SVM | 0.9703 $\pm$ 0.0018 | 0.9380 $\pm$ 0.0030 |
| AdaBoost | 0.9671 $\pm$ 0.0016 | 0.9284 $\pm$ 0.0035 |
| Logistic Regression | 0.9666 $\pm$ 0.0010 | 0.9083 $\pm$ 0.0053 |
| KNN | 0.9478 $\pm$ 0.0036 | 0.9197 $\pm$ 0.0038 |
| QDA | 0.9254 $\pm$ 0.0057 | 0.8988 $\pm$ 0.0055 |
| Native Bayes | 0.8973 $\pm$ 0.0073 | 0.8717 $\pm$ 0.0052 |

Accuracy are estimated by performing 10-fold cross-validation. Algorithms are ranked by accuracy in SNV classification. Random forest, ANN, logistic regression and KNN are set to run with eight threads. “ANN”: artificial neural network. “SVM”: single support vector machine. “KNN”: K-nearest neighbors classifier. “QDA”: quadratic discriminant analysis.

**Table S4: Variant-level quality metrics of variants in the BP dataset processed by ForestQC with different settings**

| Metric | ForestQC | ForestQC (ME rate not used) |
| --- | --- | --- |
| Total SNVs | 22227503 | 22301653 |
| Known SNVs | 19361635 | 19401450 |
| Known SNVs (%) | 87.11% | 87.00% |
| Novel SNVs | 2865868 | 2900203 |
| Novel SNVs (%) | 12.89% | 13.00% |
| Known Ti/Tv | 2.1678 | 2.1620 |
| Novel Ti/Tv | 1.7790 | 1.7577 |
| Total indels | 2789037 | 2813369 |
| Known indels | 2237002 | 2251421 |
| Known indels (%) | 80.21% | 80.03% |
| Novel indels | 552035 | 561948 |
| Novel indels (%) | 19.79% | 19.97% |
| Multi-allelic SNVs | 77693 | 78220 |
| Multi-allelic SNVs (%) | 0.35% | 0.35% |
| Known multi-allelic SNVs | 75107 | 75378 |
| Known multi-allelic SNVs (%) | 0.39% | 0.39% |
| Singletons in SNVs | 3801389 | 3804176 |
| Singletons in SNVs (%) | 17.10% | 17.06% |
| Singletons in indels | 433222 | 433035 |
| Singletons in indels (%) | 15.53% | 15.39% |

There are 20 metrics in total, which are described in Material and Methods section in detail. “Known” stands for variants in dbSNP. “Novel” stands for variants not in dbSNP. The version of dbSNP is 150.

**Table S5: Variant-level quality metrics of good variants in the BP dataset processed by different methods**

| Metric | No QC | ABHet | VQSR | ForestQC |
| --- | --- | --- | --- | --- |
| Total SNVs | 25081636 | 22415368 | 24239357 | 22227503 |
| Known SNVs | 21165051 | 19665276 | 20675746 | 19361635 |
| Known SNVs (%) | 84.38% | 87.73% | 85.30% | 87.11% |
| Novel SNVs | 3916585 | 2750092 | 3563611 | 2865868 |
| Novel SNVs (%) | 15.62% | 12.27% | 14.70% | 12.89% |
| Known Ti/Tv | 2.0751 | 2.1541 | 2.1161 | 2.1678 |
| Novel Ti/Tv | 1.5262 | 1.7987 | 1.6182 | 1.7790 |
| Total indels | 3976710 | 2670647 | 3212886 | 2789037 |
| Known indels | 3094271 | 2188996 | 2758783 | 2237002 |
| Known indels (%) | 77.81% | 81.97% | 85.87% | 80.21% |
| Novel indels | 882439 | 481651 | 454103 | 552035 |
| Novel indels (%) | 22.19% | 18.03% | 14.13% | 19.79% |
| Multi-allelic SNVs | 153836 | 26549 | 128894 | 77693 |
| Multi-allelic SNVs (%) | 0.61% | 0.12% | 0.53% | 0.35% |
| Known multi-allelic SNVs | 134108 | 24880 | 116698 | 75107 |
| Known multi-allelic SNVs (%) | 0.63% | 0.13% | 0.56% | 0.39% |
| Singletons in SNVs | 3983906 | 3650571 | 3958584 | 3801389 |
| Singletons in SNVs (%) | 15.88% | 16.29% | 16.33% | 17.10% |
| Singletons in indels | 485453 | 395798 | 325532 | 433222 |
| Singletons in indels (%) | 12.21% | 14.82% | 10.13% | 15.53% |

Four methods are compared, including no QC applied, ABHet approach, VQSR and ForestQC. There are 20 metrics in total, which are described in Material and Methods section in detail. “Known” stands for variants found in dbSNP. “Novel” stands for variants not found in dbSNP. The version of dbSNP is 150.

**Table S6: Rare variants and common variants in the BP dataset processed by different methods**

| Method | Rare SNVs | Common SNVs | Rare indels | Common indels |
| --- | --- | --- | --- | --- |
| No QC | 16304019 (65.00%) | 8777577  (35.00%) | 2219301 (55.81%) | 1757402 (44.19%) |
| ABHet | 14682507 (65.50%) | 7732821  (34.50%) | 1655360 (61.98%) | 1015280 (38.02%) |
| VQSR | 15908575 (65.63%) | 8330751  (34.37%) | 1667942 (51.91%) | 1544939 (48.09%) |
| ForestQC | 14952779 (67.27%) | 7274684  (32.73%) | 1876432 (67.28%) | 912598 (32.72%) |

The number and fraction of rare variants (MAF < 0.03) and common variants (MAF $\geq$ 0.03) in all good variants identified by different methods in the BP dataset.

**Table S7: Variant-level quality metrics of good variants identified from gray variants in the BP dataset**

| Metric | No QC | ABHet | VQSR | ForestQC |
| --- | --- | --- | --- | --- |
| Total SNVs | 2,733170 | 620497 | 2164223 | 979361 |
| Known SNVs | 1565142 | 498672 | 1281170 | 611967 |
| Known SNVs (%) | 57.26% | 80.37% | 59.20% | 62.49% |
| Novel SNVs | 1168028 | 121825 | 883053 | 367394 |
| Novel SNVs (%) | 42.74% | 19.63% | 40.80% | 37.51% |
| Known Ti/Tv | 1.3743 | 1.7730 | 1.5670 | 1.7894 |
| Novel Ti/Tv | 1.0572 | 1.1678 | 1.1256 | 1.1966 |
| Total indels | 1094234 | 194724 | 643004 | 531525 |
| Known indels | 696479 | 132387 | 540516 | 321977 |
| Known indels (%) | 63.65% | 67.99% | 84.06% | 60.58% |
| Novel indels | 397755 | 62337 | 102488 | 209548 |
| Novel indels (%) | 36.35% | 32.01% | 15.94% | 39.42% |
| Multi-allelic SNVs | 98666 | 2237 | 86457 | 61628 |
| Multi-allelic SNVs (%) | 3.61% | 0.36% | 3.99% | 6.29% |
| Known multi-allelic SNVs | 84303 | 1965 | 77772 | 59849 |
| Known multi-allelic SNVs (%) | 5.39% | 0.39% | 6.07% | 9.78% |
| Singletons in SNVs | 396426 | 72082 | 375551 | 254634 |
| Singletons in SNVs (%) | 14.50% | 11.62% | 17.35% | 26.00% |
| Singletons in indels | 107272 | 22251 | 28817 | 70447 |
| Singletons in indels (%) | 9.80% | 11.43% | 4.48% | 13.25% |

The gray variants in the BP dataset are processed by four different methods, including no QC applied, ABHet approach, VQSR and ForestQC. There are 20 metrics in total, which are described in Material and Methods section in detail. “Known” stands for variants found in dbSNP. “Novel” stands for variants not found in dbSNP. The version of dbSNP is 150.

**Table S8: Variant-level quality metrics of good variants in the PSP dataset processed by four different methods**

| Metric | No QC | ABHet | VQSR | ForestQC |
| --- | --- | --- | --- | --- |
| Total SNVs | 33273111 | 29771182 | 31281620 | 29352329 |
| Known SNVs | 25960464 | 24142744 | 24910728 | 23514257 |
| Known SNVs (%) | 78.02% | 81.09% | 79.63% | 80.11% |
| Novel SNVs | 7312647 | 5628438 | 6370892 | 5838072 |
| Novel SNVs (%) | 21.98% | 18.91% | 20.37% | 19.89% |
| Known Ti / Tv | 2.1202 | 2.1811 | 2.1643 | 2.1990 |
| Novel Ti / Tv | 1.6253 | 1.8236 | 1.7089 | 1.8131 |
| Total indels | 5093443 | 3311136 | 3682319 | 3418242 |
| Known indels | 3679990 | 2532899 | 3012662 | 2567879 |
| Known indels (%) | 72.25% | 76.50% | 81.81% | 75.12% |
| Novel indels | 1413453 | 778237 | 669657 | 850363 |
| Novel indels (%) | 27.75% | 23.50% | 18.19% | 24.88% |
| Multiallelic SNVs | 250418 | 6685 | 188180 | 146247 |
| Multiallelic SNVs (%) | 0.75% | 0.02% | 0.60% | 0.50% |
| Known multiallelic SNVs | 219411 | 6210 | 174818 | 138890 |
| Known multiallelic SNVs (%) | 0.85% | 0.03% | 0.70% | 0.59% |
| Singletons in SNVs | 14768613 | 13849361 | 14390515 | 14055519 |
| Singletons in SNVs (%) | 44.39% | 46.52% | 46.00% | 47.89% |
| Singletons in indels | 1582090 | 1350433 | 1242934 | 1393736 |
| Singletons in indels (%) | 31.06% | 40.78% | 33.75% | 40.77% |

Four methods are compared, including no QC applied, ABHet approach, VQSR and ForestQC. There are 20 metrics in total, which are described in Material and Methods section in detail. “Known” stands for variants found in dbSNP. “Novel” stands for variants not found in dbSNP. The version of dbSNP is 150.

**Table S9: Rare variants and common variants in the PSP dataset processed by different methods**

| Method | Rare SNVs | Common SNVs | Rare indels | Common indels |
| --- | --- | --- | --- | --- |
| No QC | 24864011 (74.73%) | 8409100  (25.27%) | 3381339  (66.39%) | 1712104 (33.61%) |
| ABHet | 22005560 (75.04%) | 7321250 (24.96%) | 2337310  (72.66%) | 879481 (27.34%) |
| VQSR | 23593775 (75.42%) | 7687845 (24.58%) | 2349383  (63.80%) | 1332936 (36.20%) |
| ForestQC | 22525090 (76.74%) | 6827239 (23.26%) | 2603084  (76.15%) | 815158 (23.85%) |

The number and fraction of rare variants (MAF < 0.03) and common variants (MAF $\geq$ 0.03) in all good variants identified by different methods in PSP dataset.

**Table S10: Variant-level quality metrics of good variants identified from gray variants in the PSP dataset**

| Metric | No QC | ABHet | VQSR | ForestQC |
| --- | --- | --- | --- | --- |
| Total SNVs | 3950305 | 779868 | 2746355 | 1711698 |
| Known SNVs | 2270658 | 655148 | 1748918 | 1103937 |
| Known SNVs (%) | 57.48% | 84.01% | 63.68% | 64.49% |
| Novel SNVs | 1679647 | 124720 | 997437 | 607761 |
| Novel SNVs (%) | 42.52% | 15.99% | 36.32% | 35.51% |
| Known Ti/Tv | 1.6060 | 1.8441 | 1.7801 | 1.9336 |
| Novel Ti/Tv | 1.1726 | 1.2524 | 1.2178 | 1.4236 |
| Total indels | 1596418 | 247622 | 819608 | 718606 |
| Known indels | 1009966 | 170389 | 687948 | 439129 |
| Known indels (%) | 63.26% | 68.81% | 83.94% | 61.11% |
| Novel indels | 586452 | 77233 | 131660 | 279477 |
| Novel indels (%) | 36.74% | 31.19% | 16.06% | 38.89% |
| Multi-allelic SNVs | 198530 | 541 | 165307 | 144688 |
| Multi-allelic SNVs (%) | 5.03% | 0.07% | 6.02% | 8.45% |
| Known multi-allelic SNVs | 174058 | 491 | 154111 | 137447 |
| Known multi-allelic SNVs (%) | 7.67% | 0.07% | 8.81% | 12.45% |
| Singletons in SNVs | 1276305 | 136018 | 1053352 | 715489 |
| Singletons in SNVs (%) | 32.31% | 17.44% | 38.35% | 41.80% |
| Singletons in indels | 300870 | 51339 | 139668 | 178152 |
| Singletons in indels (%) | 18.85% | 20.73% | 17.04% | 24.79% |

The gray variants in the PSP dataset are processed by four different methods, including no QC applied, ABHet approach, VQSR and ForestQC. There are 20 metrics in total, which are described in Material and Methods section in detail. “Known” stands for variants found in dbSNP. “Novel” stands for variants not found in dbSNP. The version of dbSNP is 150.

**Table S11: Running time of ForestQC and VQSR in two datasets, measured in real time**

| Method | BP SNV | BP indel | PSP SNV | PSP indel |
| --- | --- | --- | --- | --- |
| ForestQC | 17.00 min | 3.74 min | 23.24 min | 5.82 min |
| VQSR | 6.03 h | 1.21 h | 8.30 h | 1.44 h |

**Table S12: Definitions of 23 metrics for sequencing quality control calculated for sample-level and variant-level**

| Metric | Definition | Sample-level  or Variant-level |
| --- | --- | --- |
| Het / Hom | (# of heterozygous calls) / (# of homozygous non-reference calls) | Sample-level only |
| Total SNVs | The total number of SNVs | Both |
| Total SNVs (%) | The proportion of the total SNVs of a sample in the total SNVs of the entire dataset | Sample-level only |
| Known SNVs | The number of SNVs found in dbSNP | Both |
| Known SNVs (%) | The proportion of SNVs found in dbSNP | Both |
| Novel SNVs | The number of SNVs not found in dbSNP | Both |
| Novel SNVs (%) | The proportion of SNVs not found in dbSNP | Both |
| Known Ti/Tv | The Ti/Tv ratio of the known SNVs | Both |
| Novel Ti/Tv | The Ti/Tv ratio of the novel SNVs | Both |
| Total indels | The total number of indels | Both |
| Total indels (%) | The proportion of the total indels of a sample in the total indels of the entire dataset | Sample-level only |
| Known indels | The number of indels found in dbSNP | Both |
| Known indels (%) | The proportion of indels found in dbSNP | Both |
| Novel indels | The number of indels not found in dbSNP | Both |
| Novel indels (%) | The proportion of indels not found in dbSNP | Both |
| Multi-allelic SNVs | The number of multi-allelic SNVs | Variant-level only |
| Multi-allelic SNVs (%) | The proportion of multi-allelic SNVs | Variant-level only |
| Known multi-allelic SNVs | The number of multi-allelic SNVs found in dbSNP | Both |
| Known multi-allelic SNVs (%) | The proportion of multi-allelic SNVs found in dbSNP | Both |
| Singletons in SNVs | The number of singletons in SNVs | Both |
| Singletons in SNVs (%) | The proportion of singletons in SNVs | Both |
| Singletons in indels | The number of singletons in indels | Both |
| Singletons in indels (%) | The proportion of singletons in indels | Both |

Only three metrics, (Het / Hom, % Total SNVs and % total indels) are only calculated for sample-level. Other metrics are measured for every variant site and every sample. The version of dbSNP used in this study is 150.
